# RNA-seq, bioinformatic identification of potential MicroRNA-Like Small RNAs in the edible mushroom *Agaricus bisporus* and experimental approach for their validation

**DOI:** 10.1101/2021.10.13.464216

**Authors:** Francisco R. Marin, Alberto Dávalos, Dylan Kiltschewskij-Brown, Maria C. Crespo, Murray Cairns, Eduardo Andres-Leon, Cristina Soler-Rivas

## Abstract

Although genomes from many edible mushrooms are sequenced, studies on fungal miRNAs are scarce. Most of the bioinformatic tools are designed for plants or animals but fungal miRNAs processing and expression share similarities and differences with both kingdoms. Moreover, since mushroom species such as *Agaricus bisporus* (white button mushroom) are frequently consumed as food, controversial discussions are still evaluating whether their miRNAs might or might not be assimilated, perhaps within extracellular vesicles (i.e exosomes). Therefore, the *A. bisporus* RNA-seq was studied in order to identify potential *de novo* miRNA-like small RNAs (milRNAs) that might allow their later detection in the diet. Results pointed to 1 already known and 37 *de novo* milRNAss. Three milRNAss were selected for RT-qPCR experiments. Precursors and mature milRNAs were found in the edible parts (caps and stipes) validating the predictions carried out *in silico*. When their potential gene targets were investigated, results pointed that mostly were involved in primary and secondary metabolic regulation. However, when human transcriptome is used as target, the results suggest that they might interfere with important biological processes related with cancer, infectious process and neurodegenerative diseases.

## INTRODUCTION

Small RNAs (sRNAs) are a ubiquitous class of non-coding RNAs, with an average size of 20-30 nt, involved in RNA silencing pathways, also known as post-transcriptional gene silencing (PTGS) in plants, quelling in fungi and RNAi (RNA interference) in animals (1). Despite a heterogeneous group of minor and less studied sRNAs such as tiny non-coding RNAs (tncRNAs), trans-acting siRNAs (tasiRNAs), etc. (2, 3), microRNA (miRNAss) are, together with short interfering RNAs (siRNA) and PIWI-interacting RNA (piRNA), the three major categories of sRNAs. piRNAs are associated with proteins from the Piwi class of animal Argonaute (AGO) proteins, do not require RNAse III for their maturation and play important roles in germline cells. siRNAs and miRNAs are associated with the AGO proteins and share important features such as both are produced by the Dicer ribonuclease and perform their biochemical functions mainly in somatic cell lines (1, 4). However, they also show some differences; *i.e.* miRNAs are involved in the regulation of protein-coding genes through translational repression and messenger RNAs (mRNAs) degradation while siRNAs also are involved in antiviral defense. Moreover, miRNAs are originated from precursors with a typical hairpin structure generating a single miRNAs(guide):miRNAs*(passenger) duplex, while siRNAs are processed from long bimolecular RNA duplexes and a multitude of siRNAs duplexes are generated from each siRNAs precursor molecule (1, 4).

miRNAs are considered as the hallmark of the RNA silencing pathways to guide the selective degradation of mRNAs by cleavage, translational repression, or transcriptional suppression of target (mRNAs). These molecules were noticed in a wide variety of organisms, from large DNA viruses (Epstein-Barr and herpes viruses) (5) to amoeba, brown algae, nematodes, mollusks, tunicates, sea lamprey, insects, monocots, dicots, vertebrates and also in fungi (6) and, in eukaryotes, they showed highly conserved domains. Nowadays, miRNAs studies mainly focus on animal or plant leaving those on fungal kingdom behind. Animal miRNAs are typically 22 nt in length, generated from miRNAs-encoding genes that generate single-stranded RNA precursors with characteristic hairpins structures. They need specific proteins such as drosha, dicer, argonaute, etc., together with RNA pol II for their biosynthesis and exporting out of the nucleus (4). Plant miRNAs are typically 21 nt in length and include a 2′-O-methylation in 3′end, showing then a slightly different biosynthetic pathway than animals involving other enzymes and maturing steps until they are release into the cytoplasm (7). In both cases, canonical and non-canonical biosynthetic pathways co-exist being the first one the preferential path. For instance, in metazoan, non-canonical pathways produce only 1% of the reported miRNAs (4, 8). Although quelling was previously noticed (9), the presence of miRNAs in fungi was reported for the first time in *Neurospora* spp. as miRNAs-like RNAs (milRNAs) (10). Since then, several fungal milRNAs candidates were identified. For this, deep sequencing technologies and bioinformatics tools have being used, not only in other filamentous fungi (11–15), but also in basidiomycetes such as *Ganoderma spp.* (16, 17) or *Antrodia cinnamomea* (18).

Experimental studies in *Neurospora* spp. indicated that fungal milRNAs share similarities and differences with miRNAs from other kingdoms. For instance, they share with their metazoan counterparts the origins from stem-loop RNA precursors, such as happens with milR-1, the most abundant milRNAs in *Neurospora crassa (N. crassa).* Furthermore, they both require Dicer, Ago, and QIP proteins (6, 10). However, others milRNAs are generated by similar (but not identical) pathways to those of canonical miRNAs of plants and animals. Thus, four different biosynthetic pathways produce *N. crassa* milRNAs where different combinations of common elements participate together with other ones (19) and only one type of milRNAs is produced by a dicer-independent pathway (10). Moreover, their RNA precursors are transcribed mainly from intergenic regions, they show a strong preference for uracil at their 5’ and their average length is 25 nt for the mature form although a wider range between 19 and 31 nt was reported (6).

On the other hand, an increasing number of evidences on the relevance of miRNAs in the fields of Agronomy and Food Sciences are arising. Thus, very recently, it has been pointed the potential use of miRNAs as tools to increase pest tolerance in agronomic practices (20, 21). Besides, the latest evidence shows that dietary plant miRNAs can not only be absorbed in the intestine, but also be absorbed and packaged by gastric epithelial cells and then secreted into the circulatory system. The former, leads to consider miRNAs as biologically active plant-derived in a similar way that phytochemicals and, therefore, potentially responsible of food functionality (22).

Although *Agaricus bisporus* (the white-button mushroom, *A. bisporus*) is one of the most consumed mushroom worldwide and its genome was completely sequenced (H97 variety) (23), no information was found so far describing the presence of sRNAs in its edible parts (fruiting body and stipe). Therefore, this work was aimed to identify putative milRNAs from *A. bisporus* using NGS (next generation sequencing) and bioinformatics tools followed by an experimental approach with RTq-PCR to validate theoretical milRNAs candidates. Furthermore, a prediction of potential targets on *A. bisporus* genome, together with an approach to cross-kingdom putative regulation on human, was carried out, and a functional analysis of the regulated pathways both in mushroom and in human were also done. Finally, due to our theoretical predictions we expect to give a research framework for the experimental investigation.

## MATERIALS AND METHODS

### Biological material

Fruiting bodies from *Agaricus bisporus* L. (Imbach) Fungisem H-15 were kindly offered by CTICH (Centro Tecnológico de Investigación del Champiñón de La Rioja, Autol, Spain) after cultivation under controlled conditions. Fruiting bodies from the first flush where harvested before their gills were exposed (developmental stage 2-3 according to Hammond & Nichols (24). Afterwards, they were sliced, lyophilized and ground as described by Ramírez-Anguiano et al. (25). Mushroom caps and stipes were separated and prepared as indicated for the complete fruiting bodies. Mushroom powders were stored at −20°C and in darkness until further use.

### Small RNAs deep sequencing

Small RNAs was extracted from powdered *A. bisporus* fruiting bodies using mirVanaTM miRNAs Isolation Kit (Ambion®, Life Technologies) according to manufacture’s instructions and quantified using a NanoDrop2000 (Thermofisher, Spain). Then, further cDNA libraries and sequencing were performed. Briefly, low molecular RNAs were isolated by 15% TBE-urea denaturing polyacrylamide gel electrophoresis (PAGE) and ligated to specific adaptors (AGATCGGAAGAGCACACGTCT) at 3′ends. After reverse transcription, appropriate amplification and purification, the cDNA was submitted to NGS single read 1 x 50 that was carried out using an Illumina MiSeq 2000 (Illumina®, Spain). Sequencing raw data of sRNAs was deposited at the NCBI Sequence Read Archive (SRA) under accession no. PRJNA770841.

### Data analysis of small RNA and miRNAs prediction

A straightforward and preliminary bioinformatic analysis of sRNAs data was carried out with the friendly-to-use suite miARma-Seq (26) (Figure 1.2), followed by a step by step analysis for the identification of known and *de novo* miRNAs (Figure 1.1).

**Figure 1.**
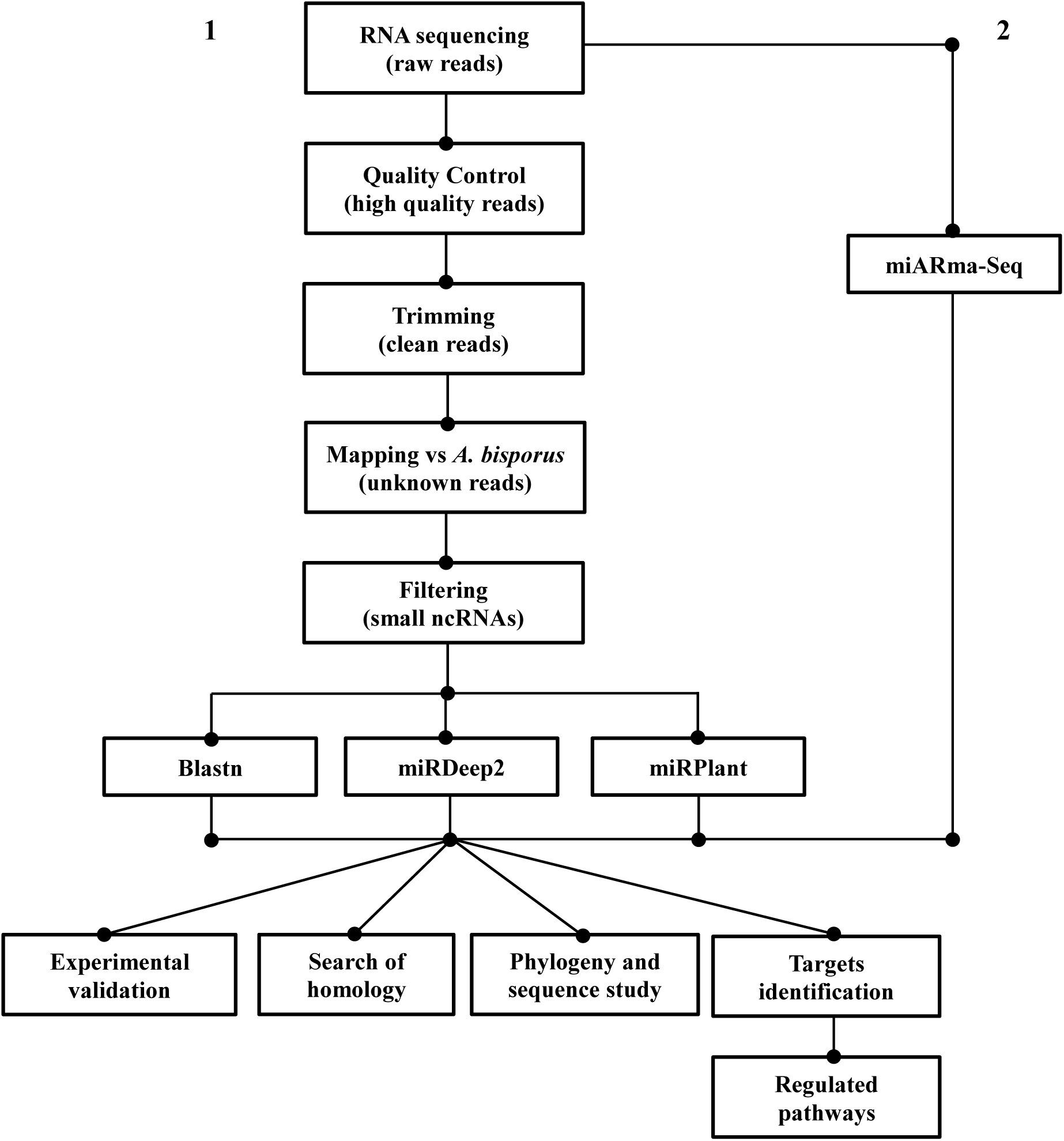
Flow-chart of the basic analysis process of milRNAs sequencing data

First, a quality control was done using FastQC (27) then, raw reads were trimmed by stripping the adaptor sequence and removing reads with lengths below 18 nt using cutadapt (28). Afterwards, clean reads were aligned against the representative genome of *A. bisporus* var. *bisporus* H97 (RefSeq assembly accession GCF_000300575.1) downloaded from NCBI (29), using bowtie 1.2.1 (30). Reads distribution within chromosomes was performed by mapping them toward *A. bisporus* var. H39 assembly (GCA_001682475.1) where the 13 scaffolds corresponded to its 13 chromosomes. Python scripts were written to complete data analysis including *i.e.* total reads counts, reads frequency etc.

The different ncRNAs (non-coding RNAs) types were determined and removed to filtered reads that might correspond to miRNAss. Thus, ncRNAs files from Ensembl Fungi (31), NCBI (32), Silva (33), GtRNAdb (34), Rfam (35) including tRNA, snRNA, snoRNA and rRNA from 80S, 60S, 5S, 5.8S, 28S, 40S, 18S, LSU, SSU eukaryote families plus the Rfam file included in miRDeep2 (36) were downloaded and eliminated from the reads by using bowtie 1.2.1.

Annotation of filtered reads with their corresponding genomic regions (*i.e.* intergenic, exonic, intronic, etc) was drawn by generating the corresponding *sam* files of perfect alignments obtained with bowtie 1. Then, they were converted into *bam* files with SAMtools (37) and from *bam* to *bed* files with BEDOPS v2.4.30 using the *bam2bed* command (38) to intersect them with the representative genome by using *intersect* from BEDTools (39). The genome in *bed* file was obtained by rearranging the *gff* format columns with *awk* (40). From the resulting file, the corresponding column was extracted with awk and regions and frequency were counted with customized python scripts.

Although miARma and miRDeep2 identify known miRNAs by comparing the reads with known sequences included in miRBase using blast (41), specific nucleotide blast (42) with filtered reads against mature and pre-miRNAs (mature.fa and hairpin.fa) from miRBase21, release 21, (43) were also carried out to improve the control of the analysis parameters. Blasts were locally run with command *blastn –task blastn-short*, a word size of 8, more suitable for miRNAs seed size (4) than by default value of 11 (44), and an e-value of 10^-4^ instead of 10, by default. Prediction of *de novo* miRNAs was carried out by miARma-seq and miRDeep2 (the latter is the central engine for miRNAs processing in miARma-Seq and therefore they both run the same algorithm) using *A. bisporus* var. *bisporus* H97 genome. An alternative prediction of *de novo* miRNAs based on criteria for plant biogenesis (instead of animal) was performed with miRPlant (45) selecting mature miRNAs length between 18 to 26 nt. In this case *A. bisporus* var. H39 assembly was used since a genome structured as chromosomes is required to run the program. Secondary structures suggested as pre-miRNAs were obtained online with RNAfold from Vienna RNA package (46).

Homologies between the predicted milRNAs in *A. bisporus* (known and *de novo*) and the sequences of 104 genomes from different Basidiomycetes species available in Ensembl Fungi (Table S1) were studied using Bowtie 1. A particular file containing most described fungal miRNAs was created to find out their potential relation with predicted *de novo* miRNAs. The file included data from *N. crassa* (10), *Fusarium oxysporum* (15), *Ganoderma lucidum* (16), *Trichoderma reesei* (13), *Penicillium marneffei* (14) and those obtained from *Pleurotus ostreatus* and *Lentinula edodes* (generated by the group but still unpublished). A multiple alignment was done by T-Coffe (47) to obtain a phylogenic tree (48) with the default values of EMBL-EBI web. Moreover, a sequence logo for milRNAs from the white button mushroom was generated with Skylign (49).

### miRNAs target prediction

The potential targets of predicted miRNAs were pointed out using psRNATarget V2, release 2017, (50) and the representative *A. bisporus* genome. Besides, a functional analysis of potential targets, on mushrooms and humans, were carried out by using the Mapper tool from Kyoto Encyclopedia of Genes and Genomes (51).

### qRT-PCR assay of miRNAs

sRNAs were extracted from mushroom caps and stipes using miRNeasy Serum/Plasma Kit (Qiagen) according to the manufacture’s protocol and quantified with a NanoDrop2000 (Thermofisher, Spain). Retrotranscription (RT) was carried out using a miScript® II RT (Qiagen) kit, then the real time PCR was used on a 7900HT Fast PCR (Applied Biosystems, Madrid, Spain) selecting 15 min at 95 °C, then 40 cycles of 15 sec at 94 °C, 30 sec at 50 °C and 45 sec at 70 °C. Samples were mixed with the SYBR Green (Qiagen) as fluorochrome. miScript Universal Primer (Qiagen) for miRNAs and primers listed in Table S2 were used in this study. Their consensus mature sequence (CMS), consensus star sequence (CSS) and consensus precursor sequence (CPS) were evaluated using rRNA 5.8S as housekeeping RNA expression with the SYBR Green real-time PCR method. An ANOVA and pair-wise Bonferroni tests were performed with R to discriminate differences between means corresponding to different samples.

## RESULTS

### Analysis of small RNA library

A total of 2,027,870 raw reads were obtained from sequencing sRNAs from *A. bisporus* fruiting bodies (Table 1). After removing 3’ specific adapters, reads showed a Q_score_ > 32 so none was discarded. Quality analysis was also carried out after trimming with identical Q_score_ results. Thus, sequences with lengths lower than 18 nt were removed following the same criteria than previous studies carried out on other fungal species (15, 16) and 1,421,021 (100%) total clean reads, corresponding to 291,880 (100%) unique reads, were obtained for further analysis. The clean reads were aligned against *A. bisporus* genome assembly and only those with perfect match were saved, resulting 1,015,249 mapped reads (71.44%) where 117,838 of them were unique reads (40.37%). After mapping, the total and unique reads length distribution in the range from 18 to 50 nt indicated that respect. 33 nt and 21 nt were the most abundant lengths within the total and unique reads (Figure 2). In the range of sRNAs (20-30 nt), 18 nt was the most abundant length in total reads and 21 nt in unique reads.

**Figure 2.**
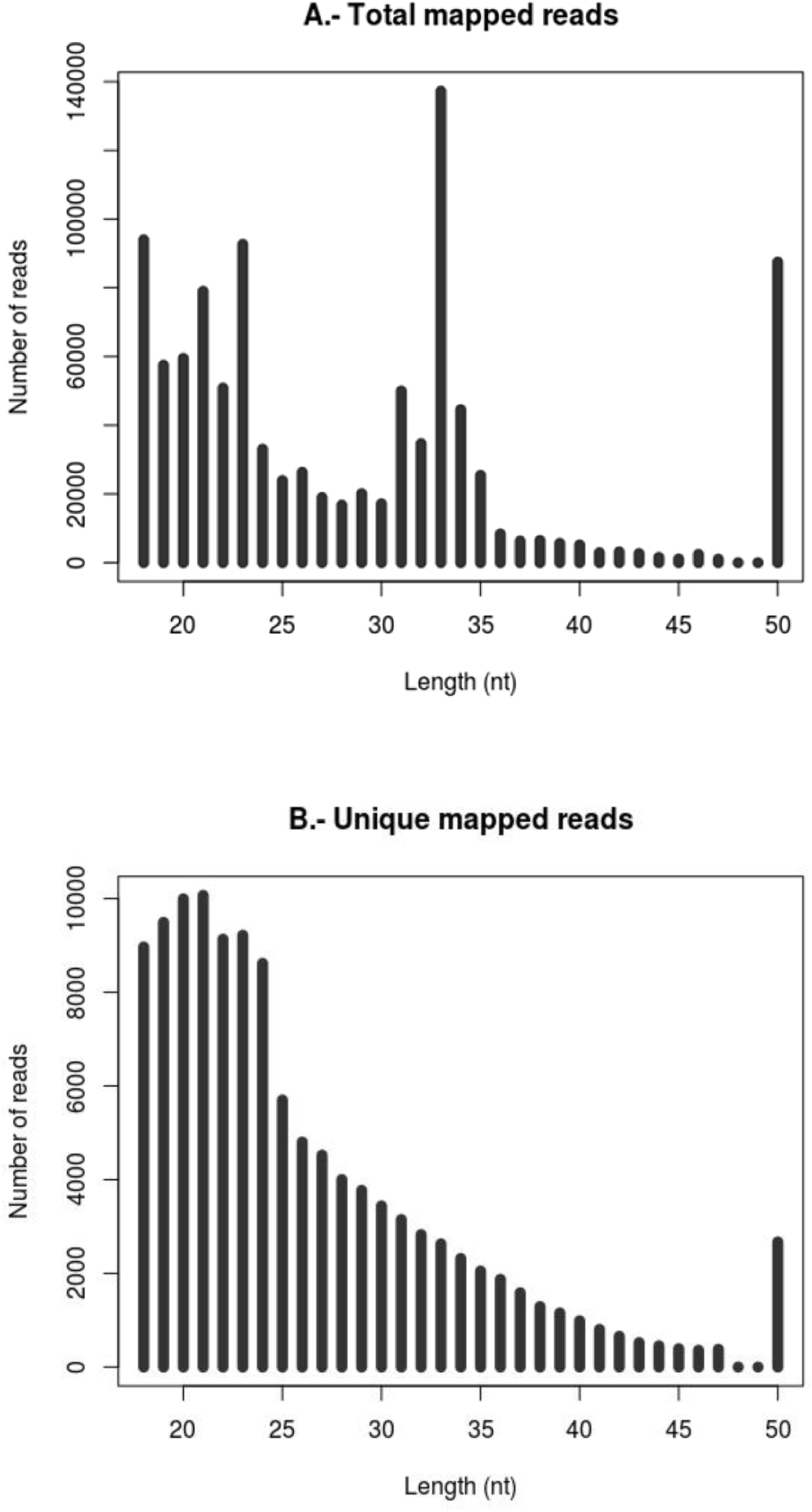
Length distribution of reads after mapping. **A.** Total mapped reads. **B.** Unique mapped reads.

**Table 1.**
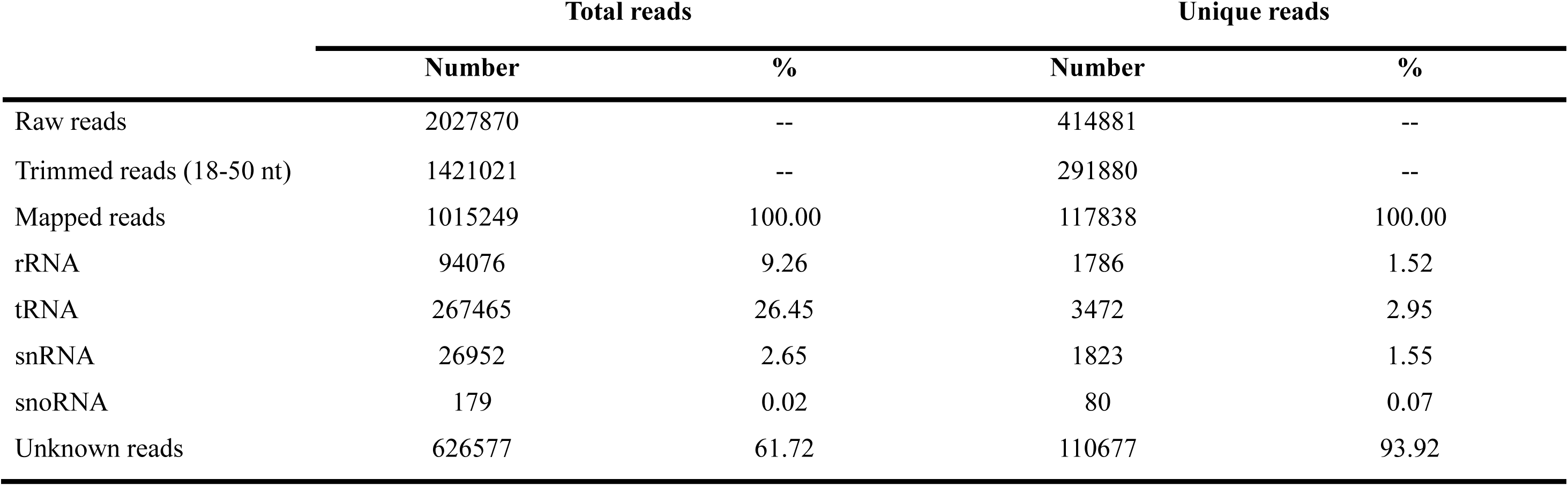
Composition of RNA library. Screening performed against an Ensembl Fungi file containing specific *A. bisporus* ncRNAs.

To avoid artifacts in miRNAs prediction, mapped reads were screened against ncRNAs from different databases (Table S3). The most restrictive results were obtained using Ensmbl Fungi files with the exception of rRNA. For this group, NCBI files showed more matchings (25.56% of total mapped reads). Although this result may lead to consider that a high percentage of the reads were originated from rRNA it is necessary to highlight that during the RNA purification on PAGE no fragmentation was carried out and therefore, RNA molecules maintained their original size (plus adaptors <50 nt) and rRNA was not isolated except for degraded fragments. Thus, since the Ensembl Fungi file was particularly designed for *A. bisporus* and the obtained results were more consistent with the experimental protocol than the other files, it was selected for filtering (and typifying) the sRNAs library (Table 1). Results indicated that a considerable amount of raw reads was sourced from tRNA (26.45% total reads) while the other ncRNAs were all together only 11.93% of total reads. Within the unique reads, only a few were ncRNAs (6.08%) indicating than most of them showed other origins (93.92%). Thus, after filtering, the 626,577 total reads and 110,677 unique reads of unknown source were mapped to figure out their chromosomal and loci distribution.

Most of the short RNA fragments (in the range 18-50 nt), before removing the ncRNAs, mapped on chromosome 9 (45% of all reads) (Figure 3A). This contribution felt to a 22% when unique reads were mapped by chromosome (Figure 3B) and to 13% when reads were filtered (ncRNAs removed) and only reads in the range 18-30 nt (sRNAs) were mapped. Considering the relatively short size of mitochondrial genome (approx.135 kbp) (52) and the large number of short RNA fragments mapped (11838 reads), mitochondrion could be pointed as an important source of sRNAs. However, after removing ncRNAs and mapping only reads in the range 18-30 nt (sRNAs) an exiguous number of 30 unique reads was found to be originated by the mitochondrion. Moreover, the unknown reads obtained after filtering were also mapped to obtain their loci annotation and results indicated that the intergenic regions were the main source of both total and unique reads representing 79.0% and 61.1% of them, respectively (Figure 4). Exonic regions accounted for only 19.9% and 37.2% of total and unique reads respectively while the contribution of intronic regions was limited to a 1.1% for total and 1.7% for unique reads.

**Figure 3.**
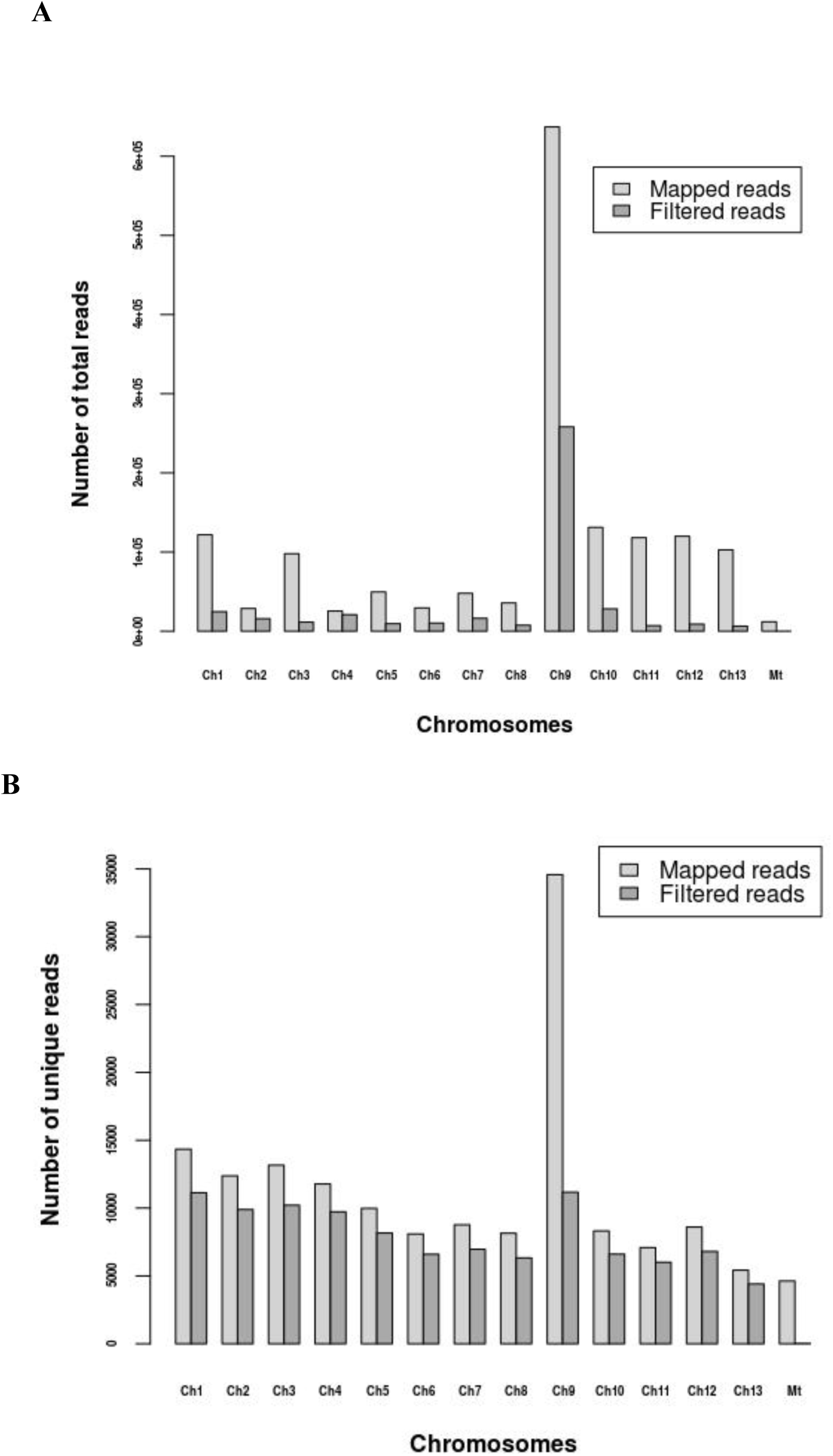
Reads distribution on chromosomes (Ch) and mitochondrion (Mt) after mapped. **A.** Distribution of total reads before and after removing ncRNAs. **B.** Distribution of unique reads before and after removing ncRNAs.

**Figure 4.**
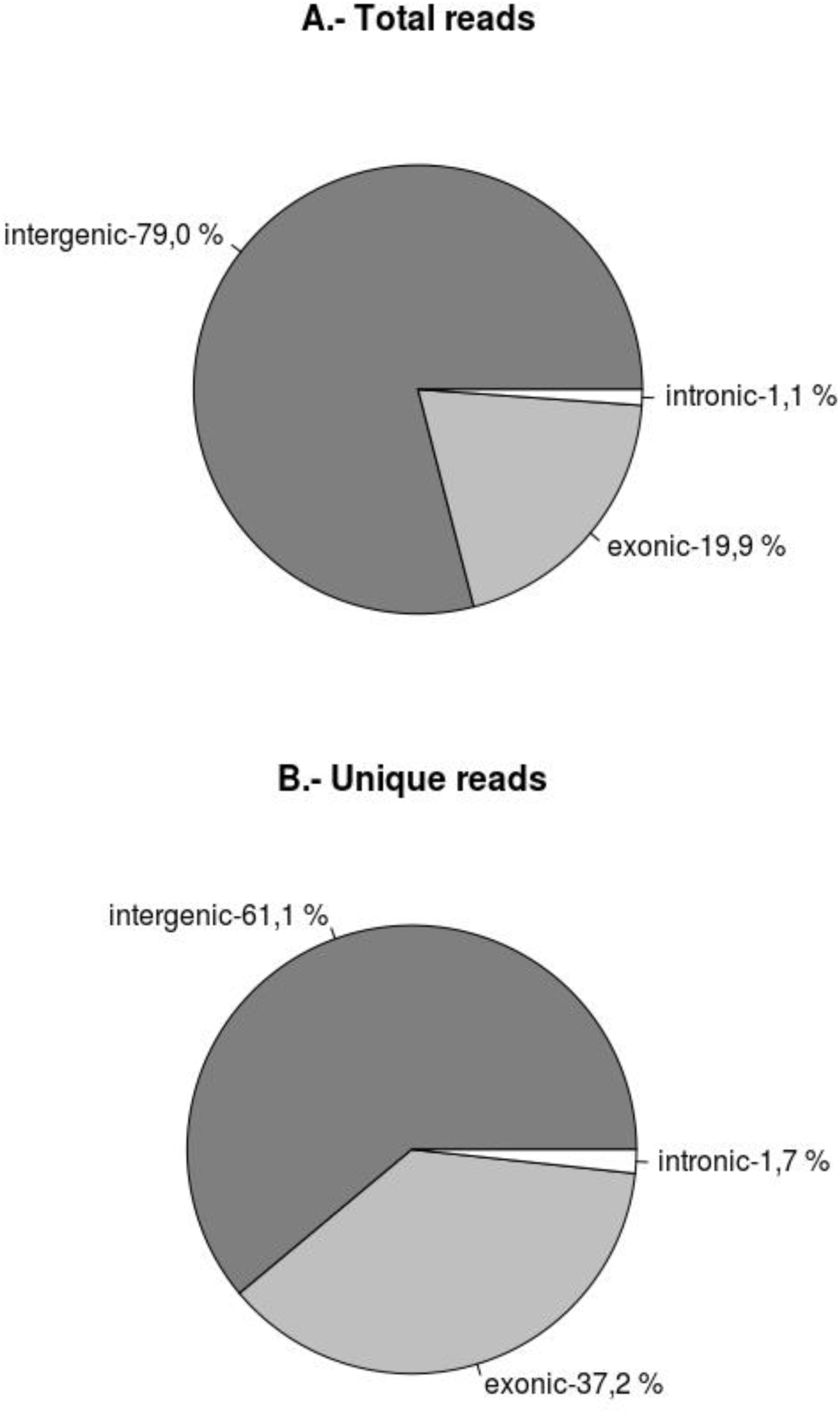
Small RNA loci annotation. Pie graphs show percentage of intergenic, exonic and intronic annotated reads. A. Total reads. B. Unique reads.

### Homology search among known miRNAs

An attempt to find homologies using miARma-Seq was performed and resulted unsuccessful. Therefore, a search against miRBase (release 21.0) was also carried out using BLAST and only three potential homologs were found among all reads. One of them matched with 67 miRNAs from different animal species and it belonged to the let-7 family but it showed only 2 reads, which led to discharge it as potential milRNAs. The other one showed homology with the plant miRNAs ptc-miR6478 and was present in higher number of reads (2060) and the third potential homolog was discarded because the match was not complete.

### Prediction of *de novo* milRNAs

Nowadays, no software is available to predict fungal milRNAs perhaps because of the scarce knowledge about them. Therefore, software designed to predict both animal and plant miRNAs was used to look for potential milRNAs in *A. bisporus*. milRNAs predictions following animal standards (with miRDeep2 and miARma-Seq) found 6 milRNAs candidates with lengths ranging from 18 to 24 nt and a mode of 21 nt and the secondary structures of their precursors could be drawn by RNAfold (Figure 5), reinforcing the theoretical prediction as contiguous sequences can generate the precursor hairpin structure (1). However, predictions following plant standards (with miRPlant) pointed a considerable higher amount of 31 milRNAs candidates, showing the same length range than miRDeep2 prediction but a mode of 22 nt. No candidates coexisted in the two predicted groups and therefore, all the potential milRNAs were listed one after the other (Table 2) and named using the nomenclature indicated in Griffiths-Jones et al. (53) and Desvignes et al. (54). The sequence logo for the predicted milRNAs suggested a tendency to include more uracil and guanine in several positions than adenine or cytosine (Figure 6). Particularly, uracil was frequent in the 5′ and 3′ extremes. The prevalence to include uracil at the 5′ extreme was previously reported for fungi from other taxons (10, 13–15) but also for the basidiomycetes such as *Ganoderma lucidum* (16).

**Figure 5.**
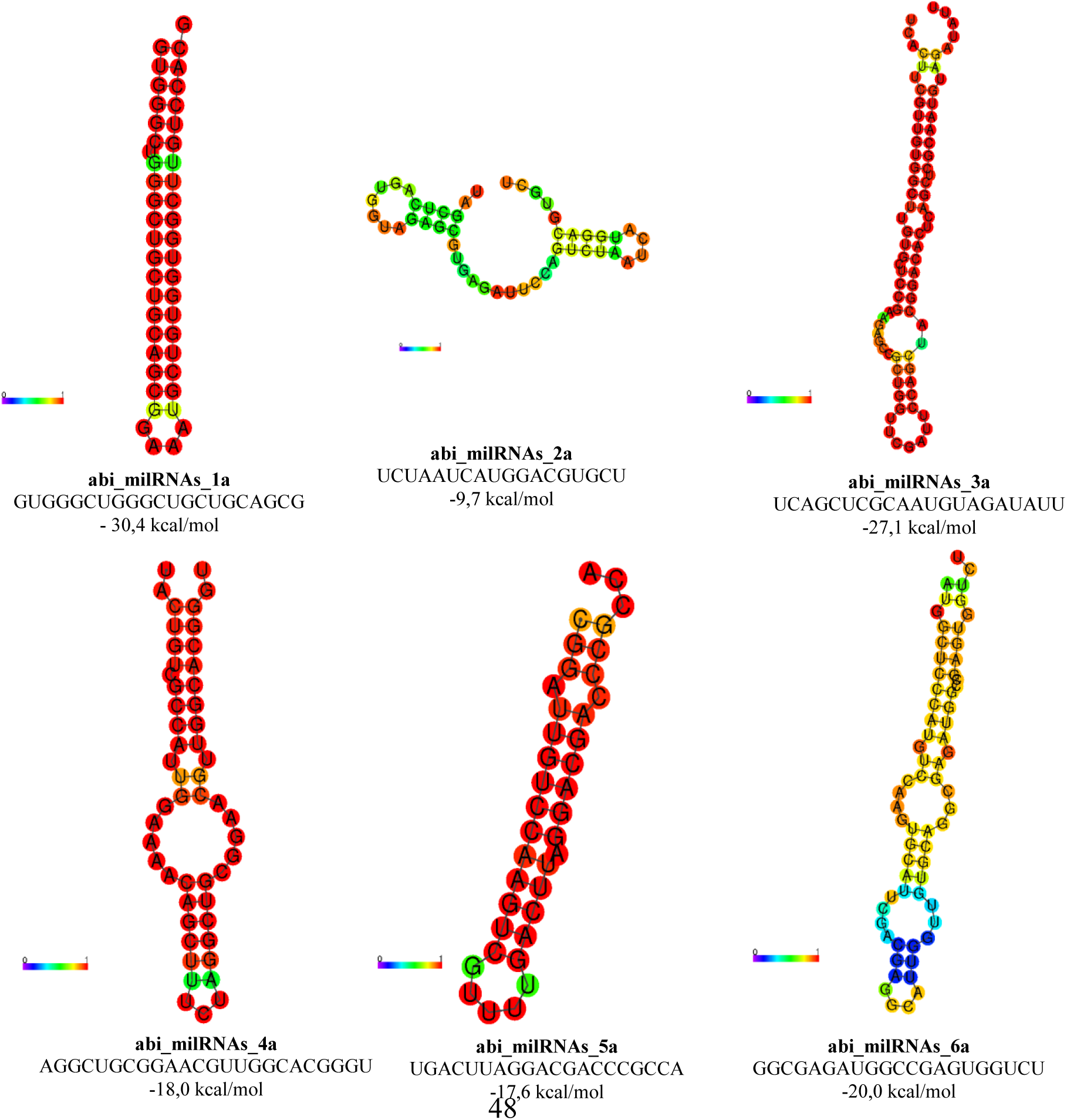
Secondary structures proposed for abi_milRNAs precursors. Colors scale (blue-red) represents the probability of each base-pair (0-1). The corresponding mature milRNAs sequence and the estimated free energy of the pre-miRNAs is also indicated.

**Figure 6.**
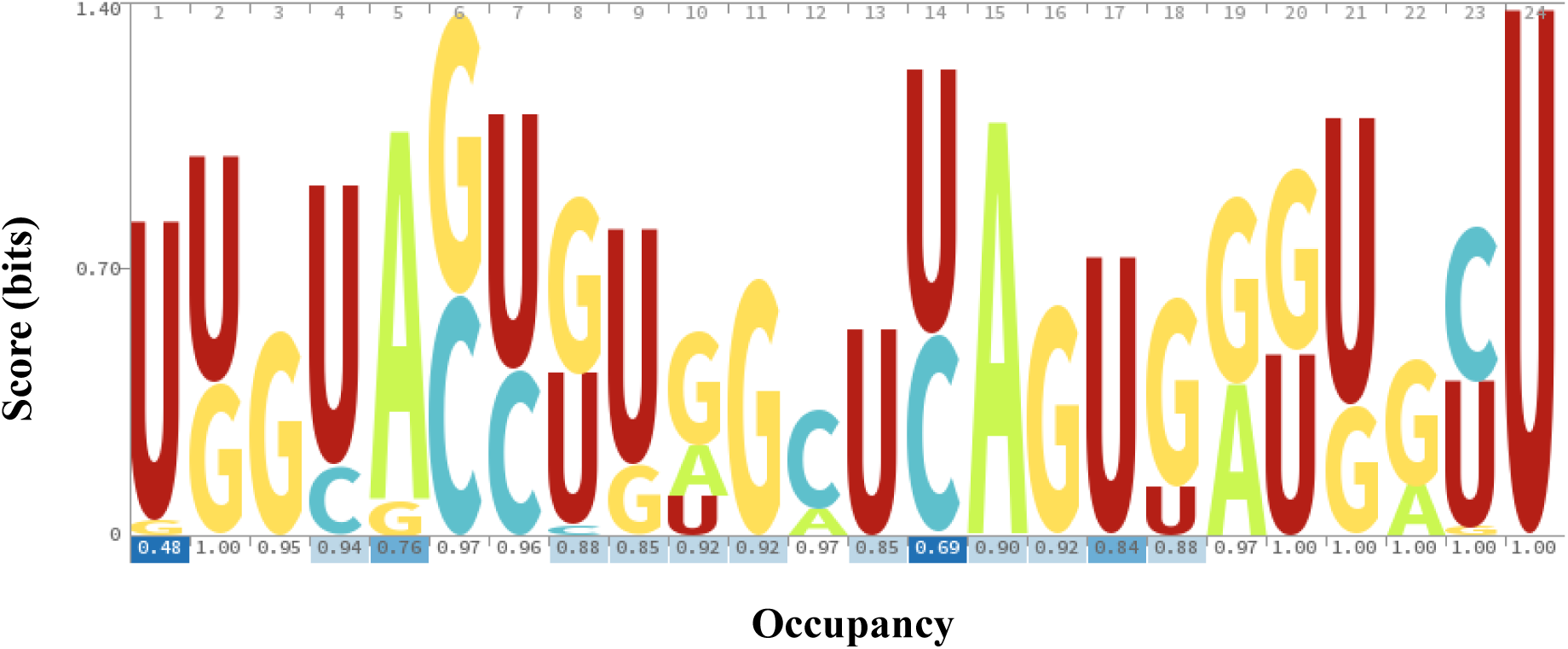
Sequence logo of predicted *A. bisporus* milRNAs. Logo represents score of weighted counts from a multiple alignment.

**Table 2.**
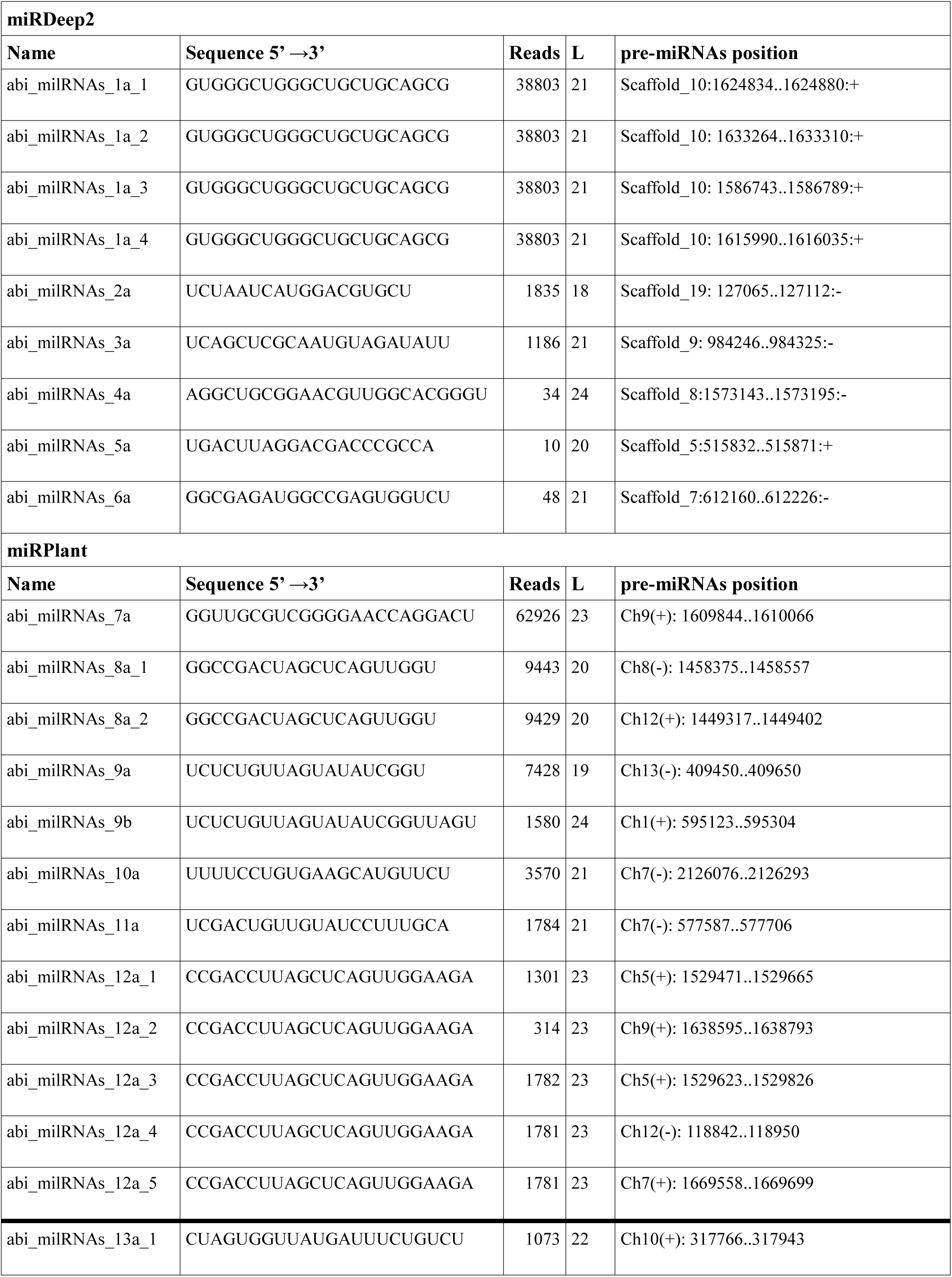
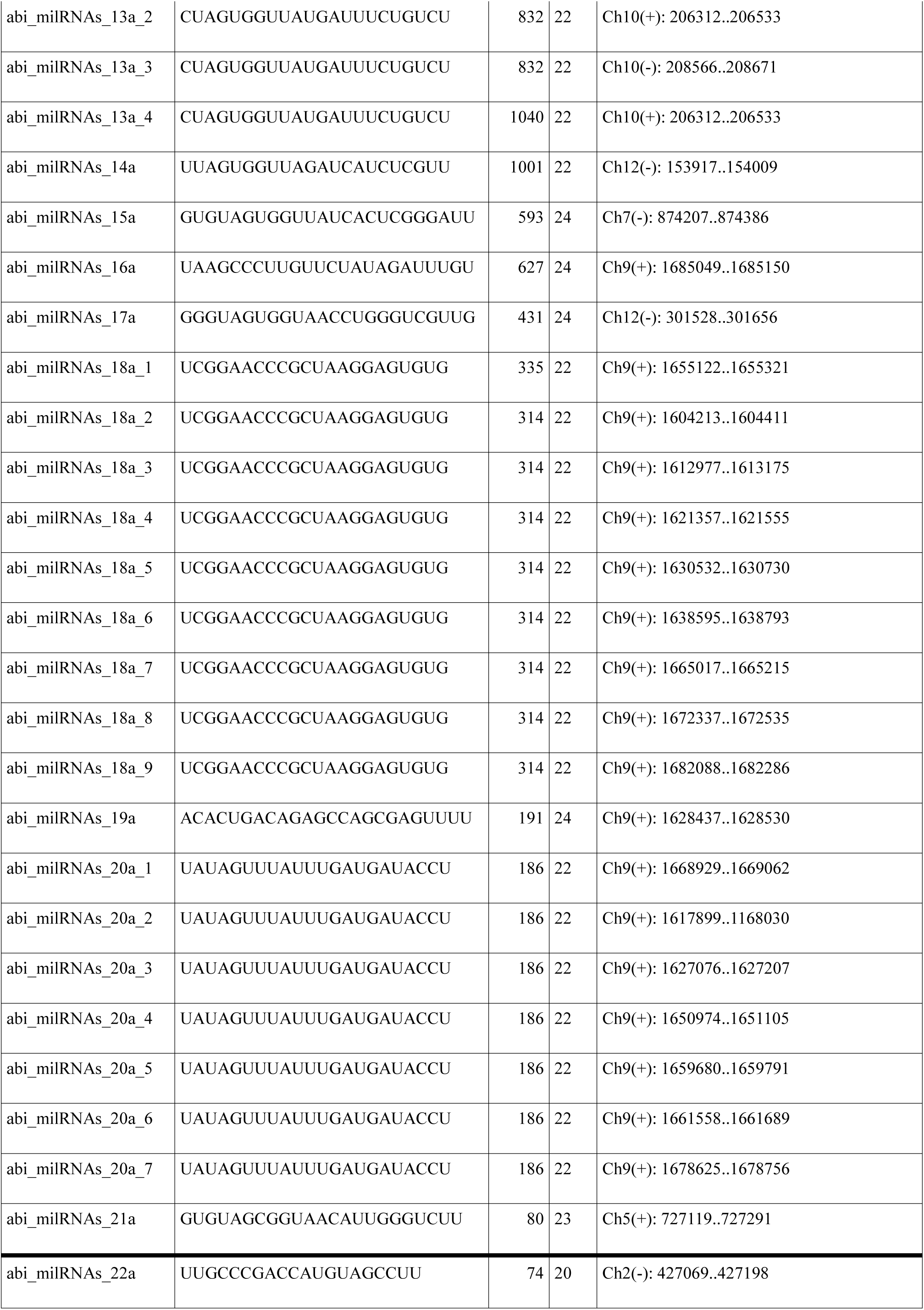
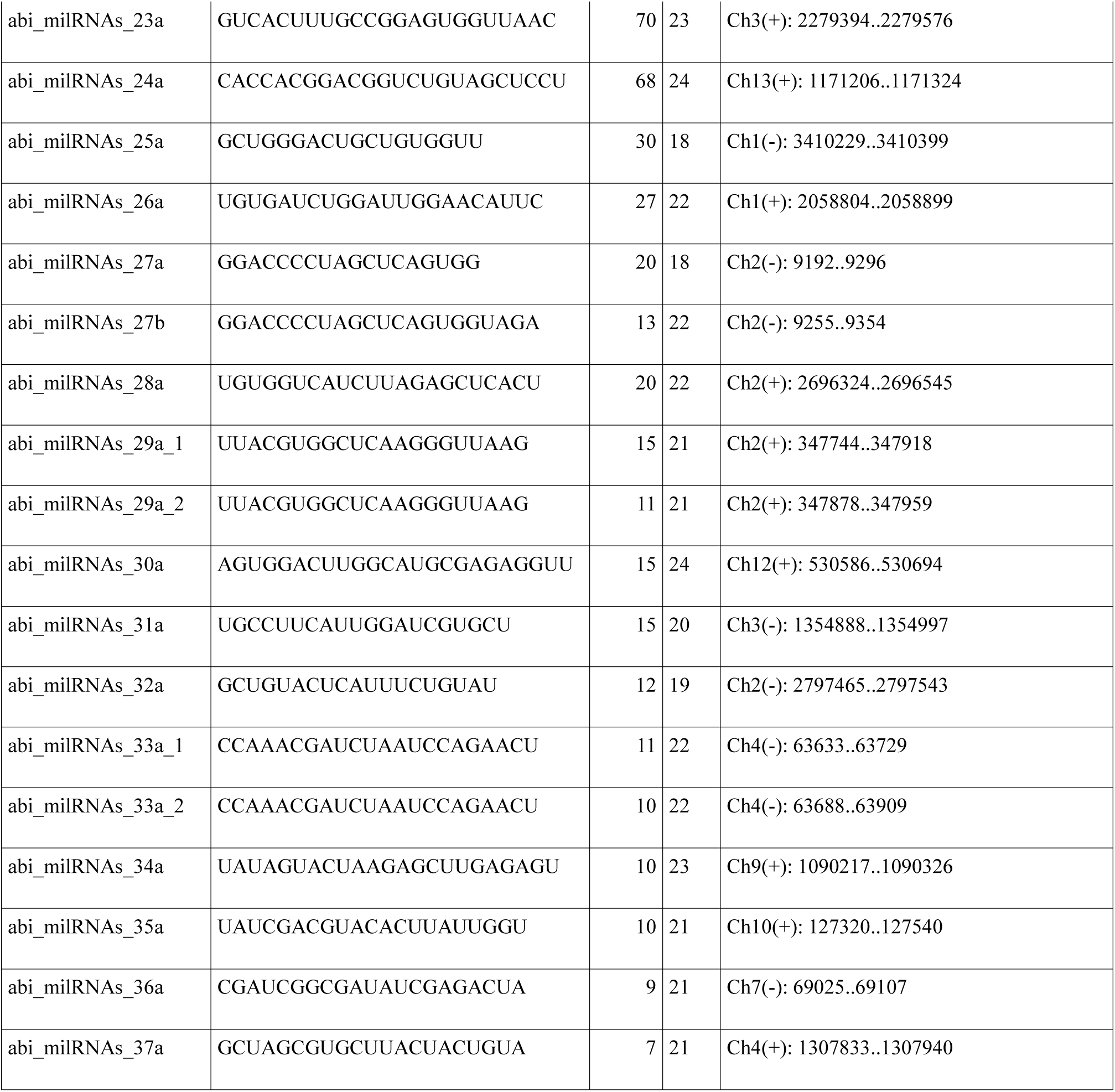
Agaricus bisporus milRNAss (Micro-Like-RNAs) candidates predicted by miRDeep2 (and miARma-Seq) and miRPlant. Reads correspond to number of reads. L: length. Pre-miRNAs position corresponds to H97 assembly for miRDeep2 group and to H39 for miRPlant one.

### Experimental verification by qRT-PCR

Three of the predicted *de novo* milRNAs (abi_milRNAs_1a, abi_milRNAs_2a and abi_milRNAs_4a), following animal criteria, were selected for experimental verification using RT-qPCR attending to their abundance. The melting curves obtained from both cap and stipe samples showed a single peak of a pure and single amplicon indicating that the designed primers achieved a proper specificity. The three predicted milRNAs were detected in both mature and precursor forms (CMS and CPS, respectively) suggesting their presence in the two mushroom tissues. The amplification curves showed CT values ranged from 8.4 to 9.7 for the qPCR to detect the mature and precursor forms of abi_milRNAs_1a, from 21.7 to 22.0 for abi_milRNAs_2a, and 33.0 to 31.0 for abi_milRNAs_4a while negative controls showed 39.0, 37.0 and 38.5 respectively. No statistically significant differential expression of pre-milRNAs was noticed between the tissues (Figure 7A). Moreover, for milRNAs expression (Figure 7B), ANOVA test did not show significant differences at α=0.05 but it did at α = 0.1. A pair-wise Bonferroni test revealed differential expression for the abi_milRNAs_4a within the cap and the stipe.

**Figure 7.**
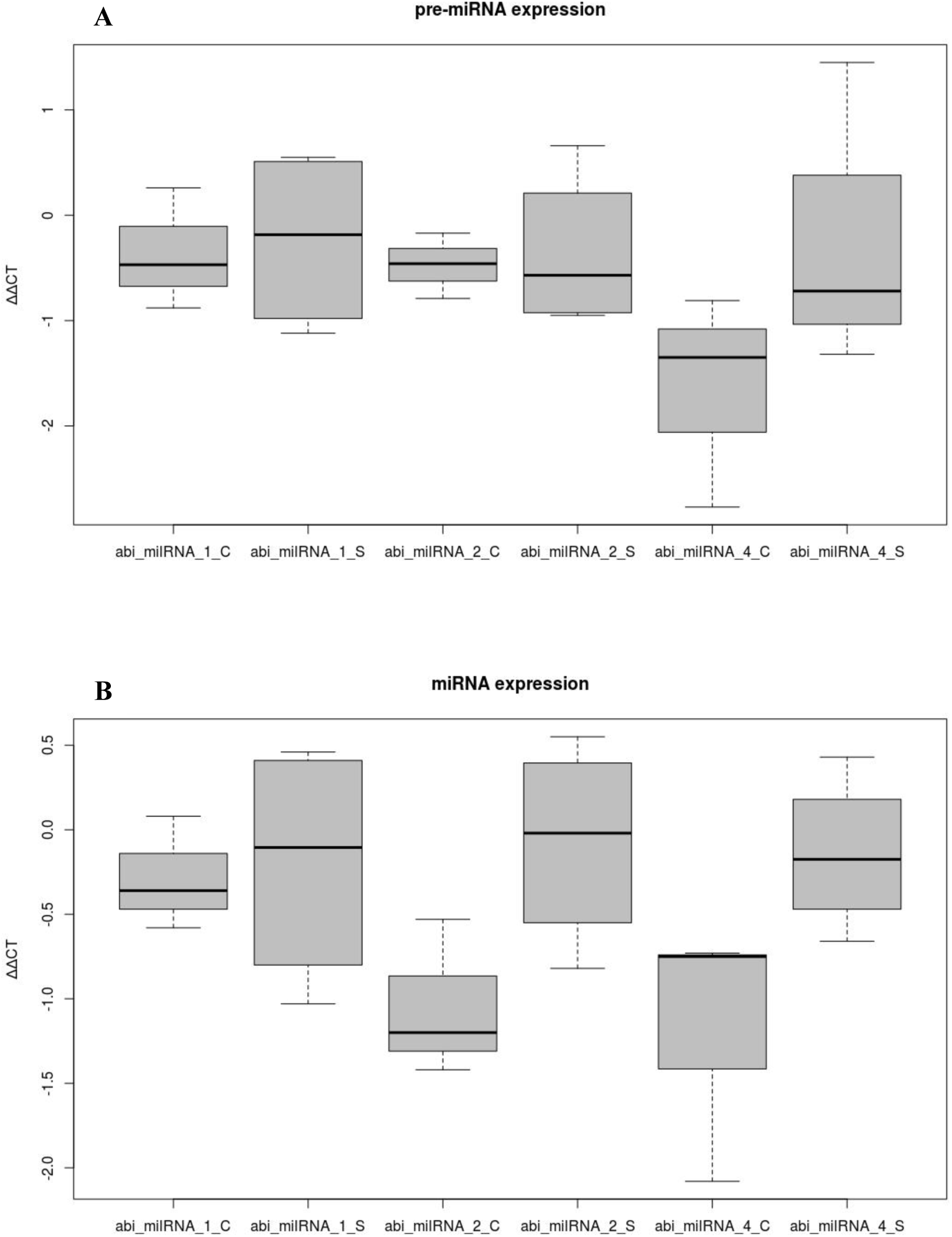
Verification of pre-miRNAs and miRNAs by RT-qPCR. **A.** Differential expression of abi_milRNAs_1a, abi_milRNAs_2a and abi_milRNAs_4a pre-miRNAs in stip (S) and cap (C). **B.** Differential expression of abi_milRNAs_1a, abi_milRNAs_2a and abi_milRNAs_4a mature miRNAs in stip (S) and cap (C).

### Homology with other fungal species

The predicted milRNAs were mapped against a collection of representative Basidiomycetes genomes (Table S4). For a conservative approach, parameters such as only 0-1 mismatch and no length variation were selected. From those, only 19 milRNAs matched with any of the studied genomes. All of them except for one (abi_milRNAs_6a) were predicted by miRPlant following biogenetic criteria for plants. The most ubiquitous *A. bisporus de novo* milRNAs were abi_milRNAs_23a, abi_milRNAs_17a, abi_milRNAs_18a and abi_milRNAs_8a since they were found in 59, 55, 52 and 52 different species, respectively, and followed by abi_milRNAs_6a (found in 49 species), predicted by miRDeep2. The predicted milRNAs perfectly matched with sequences detected in only 67 species but they showed possible homology with 94 of them when 1 mismatch was allowed. *Leucoagaricus* spp, a specie belonging to the same family that *A. bisporus* (F. *Agaricaceae*), showed the largest number of potentially identical homologues (a number of 13), followed by *Galerina marginata* and *Hebeloma cylindrosporum* with 10 identical homologues. The former mushrooms are classified in a different family (F. *Strophariaceae*) but they are included, together with *A. bisporus,* into the *Agaricales* order. In fact, the species showing 8 or 9 homologues were mostly classified in the same taxonomic order (*Agaricales*) or in the close *Polyporales* order. Only one specie, i.e. *Coniophora puteana* showed a similar number of 8 putative homologues and does not belong to *Agaricales* or *Polyporales* orders but to *Boletales* order (all belonging to the Class Agaricomycetes). When the species were included in a different taxonomical Class the number of potential homologues fall down to one or zero. On the other hand, ptc-miR6478 did not match with other fungal species.

Moreover, a file including the sequences of fungal milRNAs described in literature (up to 177 milRNAs) was created highlighting those with experimental evidences and a multiple aligning was carried out with T-Coffe to generate a preliminary phylogenetic correlation. Results indicated that several milRNAs shared internal nodes with those milRNAs with experimental evidences (Table 3). However, since the pointed species were included in a different division (*Ascomycota*) than *A. bisporus* (*Basidiomycota*), results could only suggest that the fungal milRNAs might have a common biogenesis.

**Table 3.** Fungal miRNAs with experimental evidence and their potential orthologues. (Indicated names are those given in the publication).

| miRNAs source | miRNAs with experimental evidence | miRNAs sharing an ancestor | Reference |
| --- | --- | --- | --- |
| *Agaricus bisporus* | abi_milRNAs_2a | abi_milRNAs_31a | Present work |
| *Agaricus bisporus* | abi_milRNAs_1a | abi_milRNAs_25a, pos_milRNAs_6a | Present work |
| *Fusarium oxysporum* | fox_milRNAs_2a | abi_milRNAs_6a, abi_milRNAs_17a | 15 |
| *Fusarium oxysporum* | fox_milRNAs_5 | pos_milRNAs_5a | 15 |
| *Penicillium marneffei* | PM_milR_M1 | abi_milRNAs_22a | 14 |
| *Penicillium marneffei* | PM_milR_M2 | abi_milRNAs_26a | 14 |

### Target gene prediction and functional analysis for *A. bisporus* milRNAs

*A. bisporus* milRNAs targets were studied with a theoretical approach, by using psRNATarget V2 on its reference genome and on human transcriptome. Thus, both the 37 *de novo* predicted milRNAs and ptc-miR6478 were submitted to this tool selecting default conditions. All the submitted milRNAs showed potential targets on *A. bisporus* genome, with the exception of abi_milRNAs_2a, abi_milRNAs_25a and abi_milRNAs_27a and ptc-miR6478. However, when a different software for target prediction, e.g. miRanda (55), was used under minimal restrictions (i.e. seed size and Gibbs free energy difference), targets for abi_milRNAs_2a were also found.

A total of 6946 putative targets for 35 milRNAs were predicted, being cleavage (5727 times) the most common mechanism versus translational repression (1219 times), what represents approximately 82% vs 18%. A similar pattern was followed by each individual milRNAs. Besides this, a number of 10444 genes are listed in KEGG database (56) for *A. bisporus* H97. From these, 4437 elements are regulated by the proposed milRNAs. Most of them correspond with proteins (4389) while a few ones correspond to tRNAs. After that, a functional analysis to identify the most regulated pathways was done with the KEGG-Mapper tool (51). According to the results, 109 pathways might be regulated by the proposed milRNAs, being the basic metabolic pathways, with 304 nodes, and the biosynthesis of secondary metabolites, with 124 regulated nodes, the top ones. In Table 4 are shown the 10 top regulated pathways for *A. bisporus* and human transcriptome by the proposed milRNAs. However, these results should be viewed under an annotation bias. Thus, from the list of 10444 elements, 6723 do not have KO (KEGG Orthology) identifier or a putative function assigned, i.e. we only know the function of approximately a 35% of the genetic elements of *A. bisporus*.

**Table 4.** Top 10 regulated pathways, by proposed abi_milRNAs, in *A. bisporus* and *H. sapiens* according to KEGG.

| *Agaricus bisporus* |  |  | *Homo sapiens* |  |  |
| --- | --- | --- | --- | --- | --- |
| Regulated pathway | Genes | KEGG Code | Regulated pathway | Genes | KEGG Code |
| Metabolic pathways | 304 | abv01100 | Metabolic pathways | 113 | hsa01100 |
| Biosynthesis of secondary metabolites | 124 | abv01110 | Pathways in cancer | 64 | hsa05200 |
| Biosynthesis of cofactors | 50 | abv01240 | Herpes simplex virus 1 infection | 52 | hsa05168 |
| Cell cycle | 40 | abv04111 | Pathways of neurodegeneration - multiple diseases | 47 | hsa05022 |
| Carbon metabolism | 35 | abv01200 | Human papillomavirus infection | 40 | hsa05165 |
| Autophagy | 33 | abv04138 | MAPK signaling pathway | 38 | hsa04010 |
| MAPK signaling pathway | 32 | abv04011 | PI3K - Akt signaling pathway | 35 | hsa04151 |
| Protein processing in endoplasmic reticulum | 32 | abv04141 | Proteoglycans in cancer | 33 | hsa05205 |
| Nucleocytoplasmic transport | 31 | abv03013 | Amyotrophic lateral sclerosis | 32 | hsa05014 |
| Spliceosome | 31 | abv03040 | Alzheimer disease | 32 | hsa05010 |

On the other hand, *A. bisporus* is a frequently consumed edible mushroom (57). Thus, although is necessary to clarify whether *A. bisporus* milRNAs might be absorbed with diet and reach their target in humans, we performed a theoretical approach on human transcriptome. Thus, a total of 2269 putative targets for abi_milRNAs were predicted, being cleavage (1837 times) the most common mechanism versus translational repression (432 times), what represents approximately 81% vs 19%, and a similar pattern was followed by each individual milRNAs. At the same time, and as described above for *A. bisporus* genome, abi_milRNAs_2a, abi_milRNAs_25a and abi_milRNAs_27a did not show potential targets on human transcriptome and in the same way as described for *A. bisporus*, targets for abi_milRNAs_2a were also found when a different bioinformatic tool was used. Besides this, a number of 22,214 genes are listed in KEGG database (56) for *Homo sapiens*. From these, 19,572 code for proteins and 2,642 do it for RNA elements. According to the results, 319 pathways might be regulated by the prosed *A. bisporius* milRNAs. From these, as shown in Table 4, human basic metabolic pathways, with 113 regulated nodes, and human pathways in cancer, with 64 regulated nodes, were the most regulated ones. Furthermore, it should be noticed that also 28 pathways are related to infectious process (e.g. herpes simplex virus infection, with 52 regulated nodes, and human papillomavirus infection, with 40 regulated nodes) and 7 with neurodegenerative process.

## DISCUSSION

The combination of high-throughput sequencing technologies and bioinformatics processing facilitates the identification of small RNAs in any organism, providing quantitative information about their transcription and sequence. Given that it is expected that they are highly conserved, within closely related species, and share similar biosynthetic pathways computational tools can be used to study them, *i.e.* the secondary structure of their precursor or miRNAs mature sequences, to find homologies, to screen the complete genome of multiple species, even to suggest their potential target genes among other applications. However, several of these programs were designed taking into consideration the particularities of plant or animal sRNAs and therefore, the proposed predictions for organisms from other kingdoms might be carefully look through and experimental confirmations should be always required. Therefore, for the identification of fungal milRNAs from the white-button mushroom several tools were utilized and compared and also an experimental approach was also carried out for some predicted miRNAs.

The ratios between clean reads (total reads and clean reads vs unique clean reads) obtained after sRNA sequencing were similar to the previously obtained for other fungi and the most abundant ones were those of 21 nt, after mapping for unique reads (15, 16). Other common features with fungal milRNAs were their genomic positions, being intergenic regions the major source of sRNA production (6, 15) or their strong preference for 5′U (10, 14) suggesting that they all followed a common evolutionary pathway. Furthermore, the production of sRNAs, in *A. bisporus*, seems to follow a hot spot pattern as described for other fungi (15). Thus, chromosome 9 is source of 45% of total reads and 22% of unique sequences. However, certain differences were also found when compared with less evolved fungi, such as *F. oxysporum,* where reads were mostly located in some chromosomes, 2 and 4, but also in the mitochondrion (15), which was finally irrelevant in *A. bisporus*. The low number of identified milRNAs (1 known and 37 predicted) also diverged from other basidiomycetes such as *G. lucidum* where 166 potential milRNAs were pointed (16). The latter discrepancy might be because different bioinformatics tools, and settings, were used for their identification. Indeed, miRDeep2 uses an algorithm based in a probabilistic model of animal miRNAs biogenesis that score patterns and frequencies of the RNA sequences, taking into account the suggested secondary structure (as pre-miRNAs) obtained with RNAfold from Vienna RNA package while miRPlant was similarly designed but for plant miRNAs following biogenetic plant criteria (36, 45). Although none of both tools were specifically designed for fungal milRNAs, putative candidates were postulated because apparently biogenesis of fungal milRNAs share similarities with both plant and animal kingdoms as previously mentioned (6).

On the other hand, initially we approached the search of *A. bisporus* milRNAs with an evolutionary bias, which assumes that the animal and fungal lineages share a more recent common ancestor than either does with the plant, alveolate, or stramenopile lineages (58). Thus, a prediction under biogenetic animal criteria yielded only 6 *de novo* milRNAs and experimental validation, through qPCR, was carried out for 3 of them with positive results. However, when an approach based in different biogenetic criteria was done, we found a considerably higher number of *de novo* milRNAs (31 milRNAs). Unfortunately, the shortage of funds did not allow us to validate them experimentally. However, the aforementioned features (i.e. length, hairpin precursor, thermodynamic viability –evaluated through psRNAtarget-, match with genome, homology with related species, etc.) point more to a consistent prediction, than a false one, that should be validated in further studies.

The higher number of fungal milRNAs pointed by miRPlant might suggest the hypothesis that some of the milRNAs generated by the mushroom are more closely related to plants than to animals. This observation might be in line with the fact that in the nature, many fungi establish closer environmental relations with several trees and plants forming mycorrhizas. The precise involvement of certain miRNAs in this symbiotic/parasitic nutrient exchange (beside secretion of certain signaling compounds) is not fully elucidated yet. In this sense, a different miRNAs pattern is expressed in plants with or without mycorrhizas (59, 60) and during infectious process (61). In addition, fungal milRNAs have been pointed out as key elements in this inter-kingdom talk (62) and technological approaches, also for siRNAs, to increase pest tolerance in plants have been suggested (20, 21). Therefore, it might be possible that fungal milRNAs would have needed to mimic plant miRNAs to enable a better fitting between two organisms belonging to two different kingdoms in a co-evolutionary process.

Beyond the former, and as previously mentioned, *A. bisporus* is one of the most produced mushrooms (57) and is part of our diet. To this respect, since Zhang et al. (63) reported that exogenous plant miRNAs were able to target the mammalian LDLRAP1, an increasing number of papers have been published on this topic. For instance, the early research of Baier et al (64) detected miR-29b and miR-200c, both are bovine miRNAs, in plasma after cow milk intake while Chin et al. (65) proposed a cross-kingdom inhibition of breast cancer growth by plant miR159. Additionally, accumulating evidence indicates that sRNAs can be transferred within cells and tissues and even across species (66) and miRNAs from diet have been proposed to play a role on disease prevention (67, 68). Additionally, several authors report that dietary-miRNAs can be absorbed with diet and reach their target mRNAs, both for animal (64, 69) and plant (22, 65) foodstuffs.

Although specific studies to clarify whether *A. bisporus* milRNAs could be absorbed with diet and reach their targets are needed, theoretical predictions can be useful for a non-blindly search and to narrow and guide experimental research. To this respect, while abi-milRNAs are proposed to regulate basic metabolic pathways on *A. bisporus* (Table 4) they could be involved in the regulation of several pathological process in humans. Thus, a third of the human pathways potentially regulated by abi-milRNAs are involved in diseases, i.e. 52 related to cancer, 28 to different infectious processes (including the infectious process by SARS-CoV-2: KEGG pathway hsa05171) and 7 to neurodegenerative diseases. What’s more, we find the top 10 regulated pathways (Table 4), all of them are of great relevance from a clinical perspective.

Moreover, it is wise to mention that recent publications pointed out with clinical studies the effect of specific mushrooms to prevent, for example, Alzheime’s onset (70) or improve immune status in immunocompromised breast cancer patients (71). *A. bisporus* extracts were also able to interfere human prostate cancer cells (72). To this respect, the above-mentioned diseases might potentially be regulated by abi_milRNAs according to the theoretical prediction. For instance, abi-milRNAs may regulate approximately 16% of prostate cancer nodes, according to KEGG Pathway database, leading, again to the question if dietary miRNAs are/ or not playing a biological role such as pythochemicals do.

In overall, our data here presented provide evidence that the edible mushroom *Agaricus bisporus* contain miRNA-like small RNAs, with a collection of characteristics consistent with that of fungal miRNAs. Besides, *A. bisporus* milRNAs share also characteristics with miRNAs of plant origin more than with animal miRNAs and, at least for some of the latest, experimental validation has been given. Finally, some pathways (both in mushroom and in human) are proposed to be (putatively) modulated by the presented *A. bisporus* milRNAs.

## ACKNOWLEDGMENTS

We wish to thank to CCC-UAM (“Centro de Computación Científica – UAM”) by the use of computer equipment and technical support.

## AUTHOR CONTRIBUTION

FRM and CS-R conceived the idea and supervised the work. AD, MCC, DK-B, MC and EA-E, conducted experiments. FRM, CS-R, AD and MCC wrote the paper. All authors have approved the final version of the manuscript.

## SUPLEMENTORY MATERIAL

**Table S1.**
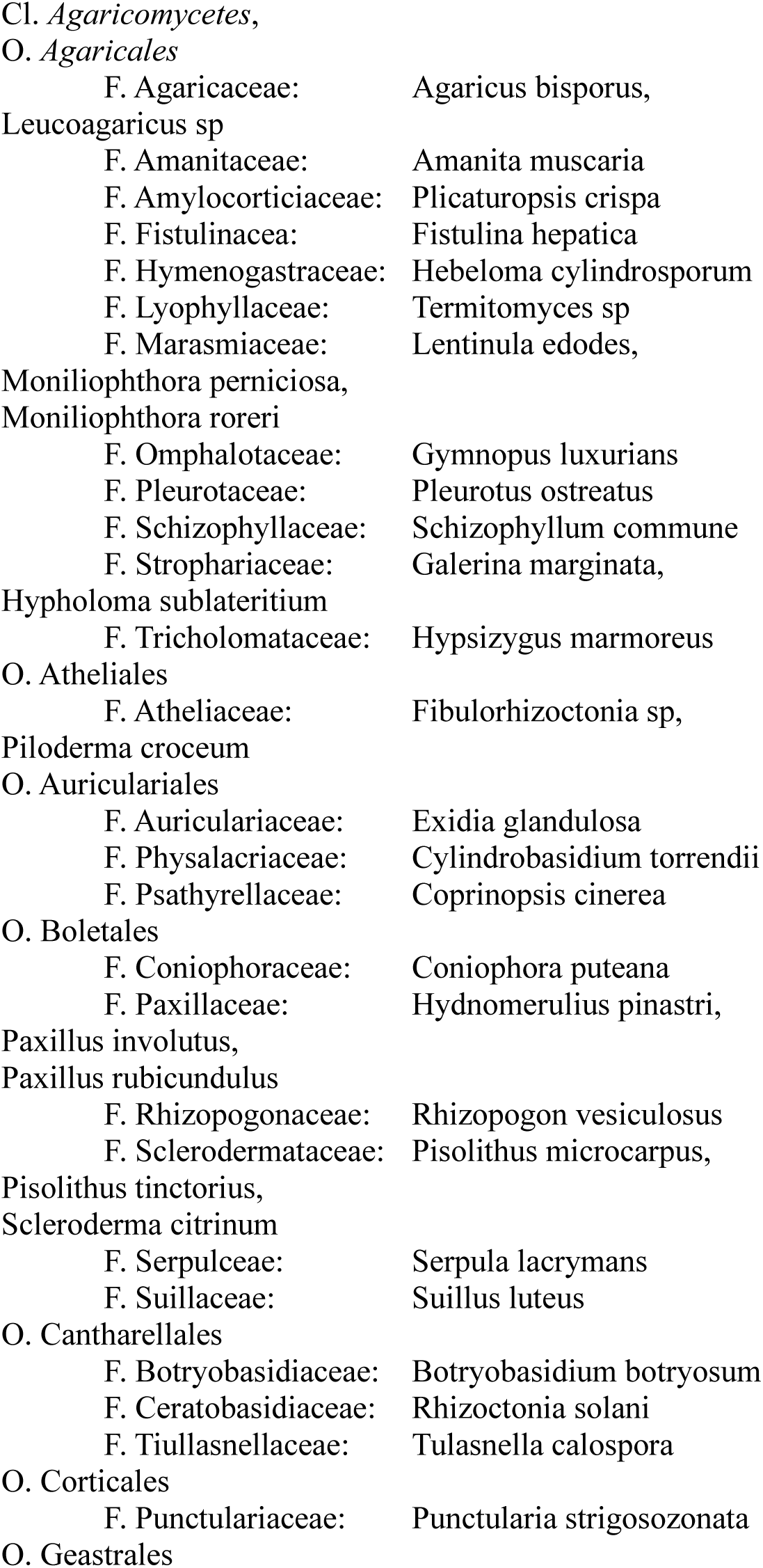
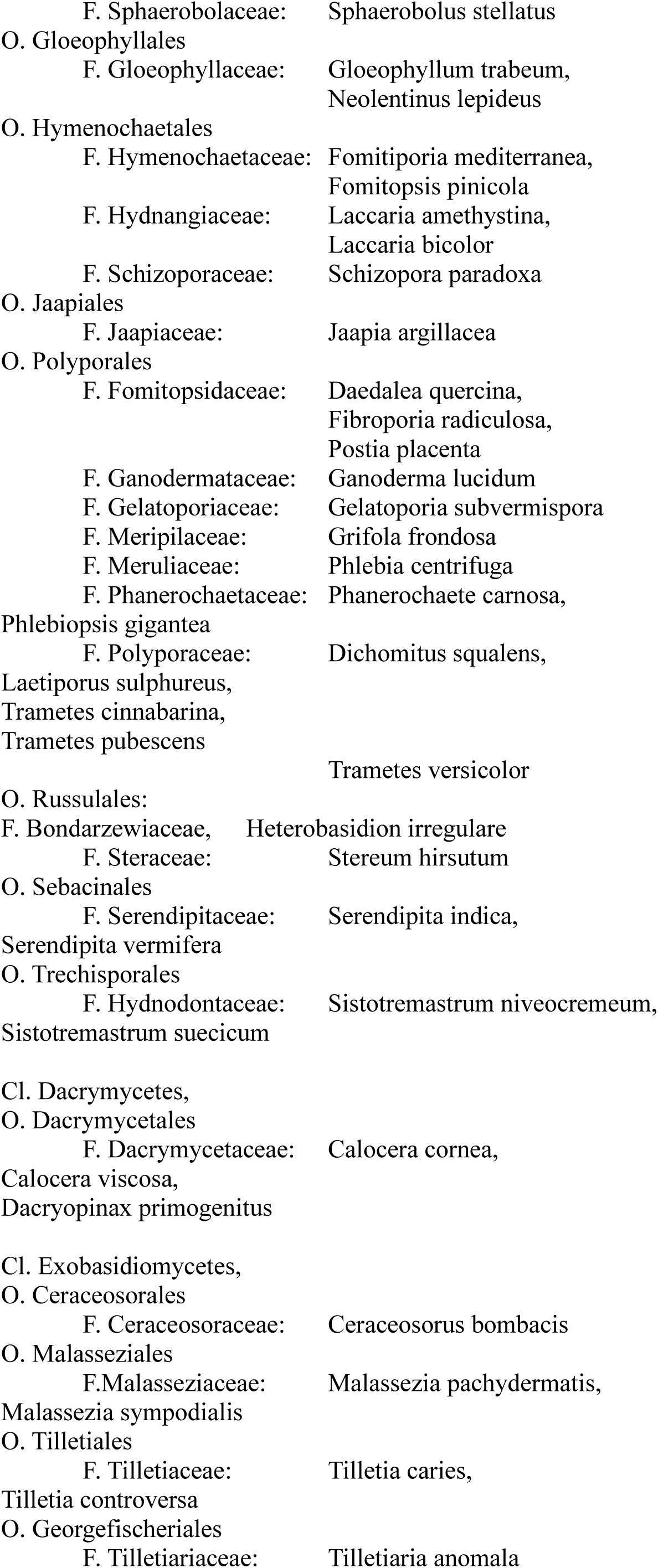
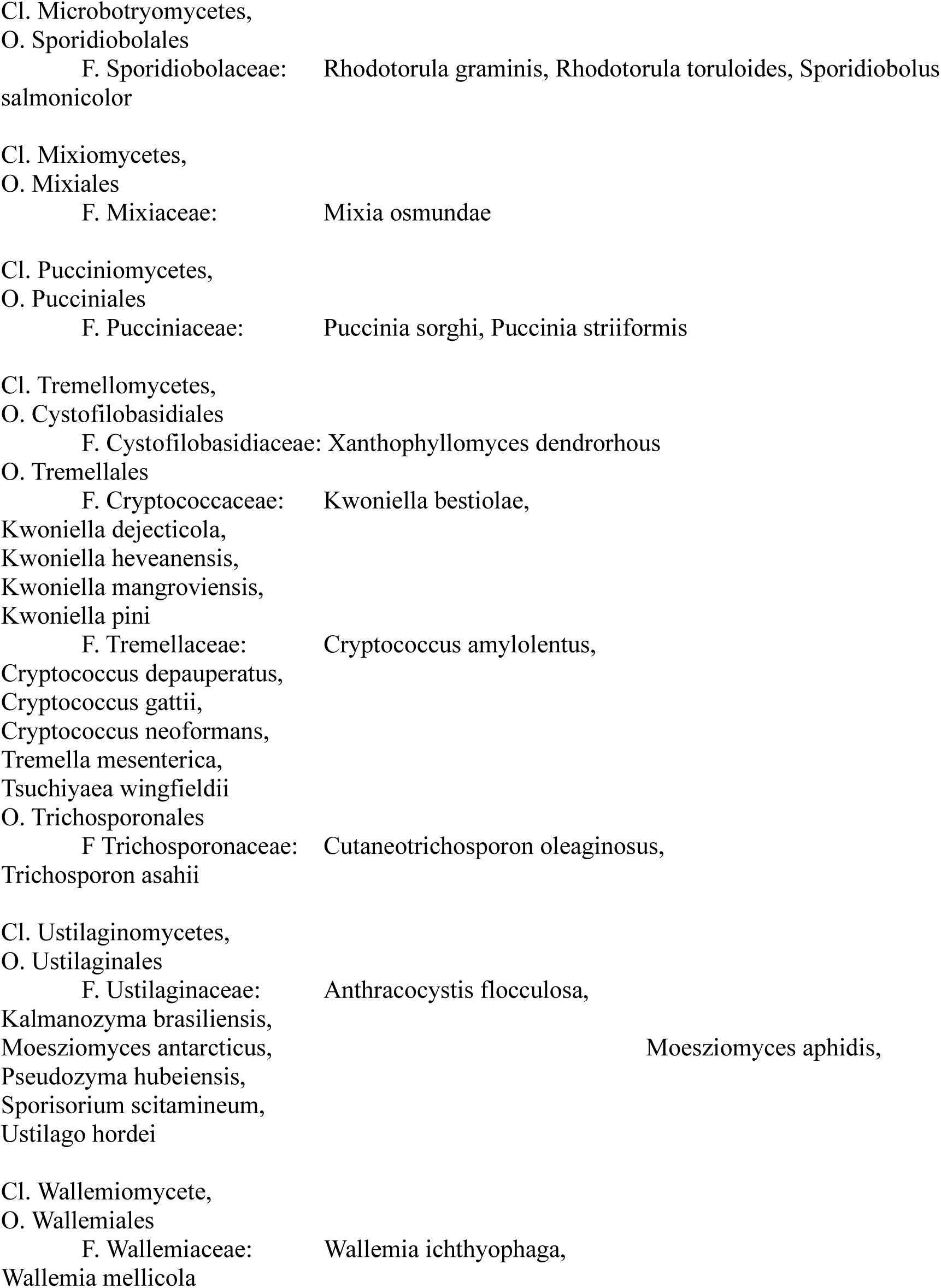
List of Basidiomycetes species downloaded for the study of homology. The species are listed following taxonomical criteria. Source: Ensembl Fungi. (ftp://ftp.ensemblgenomes.org/pub/fungi/release-37/fasta/fungi_basidiomycota1_collection/)

**Table S2.**
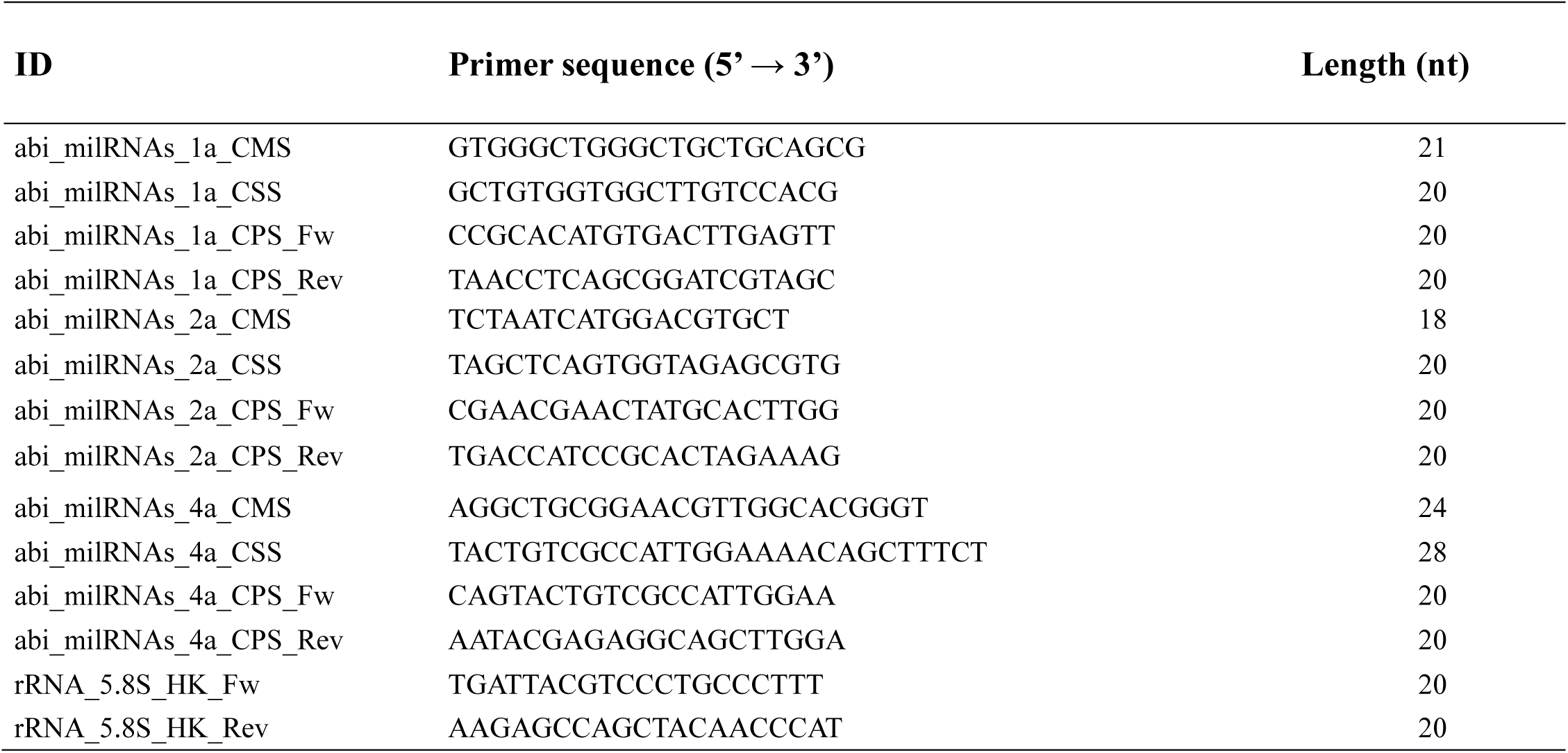
Primers used for RT-qPCR experiments in this study. CMS: consensus mature sequence; CSS: consensus star sequence; CPS: consensus precursor sequence; HK: Housekeeping; Fw: Forward; Rev: Reverse.

**Table S3.** Screening against different classes of ncRNAs from different databases. In the second column, the number between brackets indicates file features: **1.** Rfam file downloaded from miRDeep2. **2.** Rfam file corresponding to different RNA classes (i.e. snRNA, tRNA, etc) of *Agaricus bisporus*. **3.** Ensembl Fungi file corresponding to different RNA classes of *Agaricus bisporus*. **4.** NCBI-Nucleotides file corresponding to different RNA classes of *Fungi*. **5.** Raw files from other databases without refining their contents. For instance, Silva file contains all available rRNA without attending taxonomic group.

| RNA class | Database | Total reads | Unique reads |
| --- | --- | --- | --- |
| Mapped reads |  | 1015249 (100.00 %) | 117838 (100.00 %) |
| tRNA | Rfam (1) | 23968 (2.36%) | 840 (0.71 %) |
|  | Rfam (2) | 0 (0.00%) | 0 (0.00%) |
|  | Ensembl_Fungi (3) | 267465 (26.45%) | 3472 (2.94 %) |
|  | NCBI (4) | 1 (0.00%) | 1 (0.00 %) |
|  | GtRNA (5) | 1119 (0.11%) | 120 (0.10 %) |
| snRNA | Rfam (2) | 0 (0.00%) | 0 (0.00%) |
|  | Ensembl_Fungi (3) | 26952 (2.65 %) | 1823 (1.54%) |
|  | NCBI (4) | 1634 (0.16 %) | 87 (0.07 %) |
|  | SILVA (5) | 11 (0.00 %) | 8 (0.01 %) |
| snoRNA | Rfam (2) | 0 (0.00%) | 0 (0.00%) |
|  | Ensembl_Fungi (3) | 179 (0.02 %) | 80 (0.07%) |
|  | NCBI (4) | 00 (0.00%) | 0 (0.00 %) |
|  | SILVA (5) | 1 (0.00%) | 1 (0.00 %) |
| rRNA | Rfam (1) | 119153 (11.74 %) | 4217 (3.59 %) |
|  | Rfam (2) | 13 (0.00 %) | 11 (0.01 %) |
|  | Ensembl_Fungi (3) | 94076 (9.26 %) | 1786 (1.52%) |
|  | NCBI (4) | 259581 (25.56 %) | 18174 (15.42%) |
|  | SILVA (5) | 55 (0.01%) | 8 (0.01 %) |
| ncRNA | miRDeep (1) | 136120 (1.29 %) | 5057 (4.29%) |
|  | Rfam (2) | 13 (0.00 %) | 11 (0.01) |
|  | Ensembl_Fungi (3) | 388675 (38.42 %) | 7164 (6.08 %) |
|  | NCBI (4) | 261201 (25.72 %) | 18253 (15.49 %) |
| lncRNA | NONCODE (5) | 109 (0.01 %) | 84 (0.07 %) |

**Table S4.**
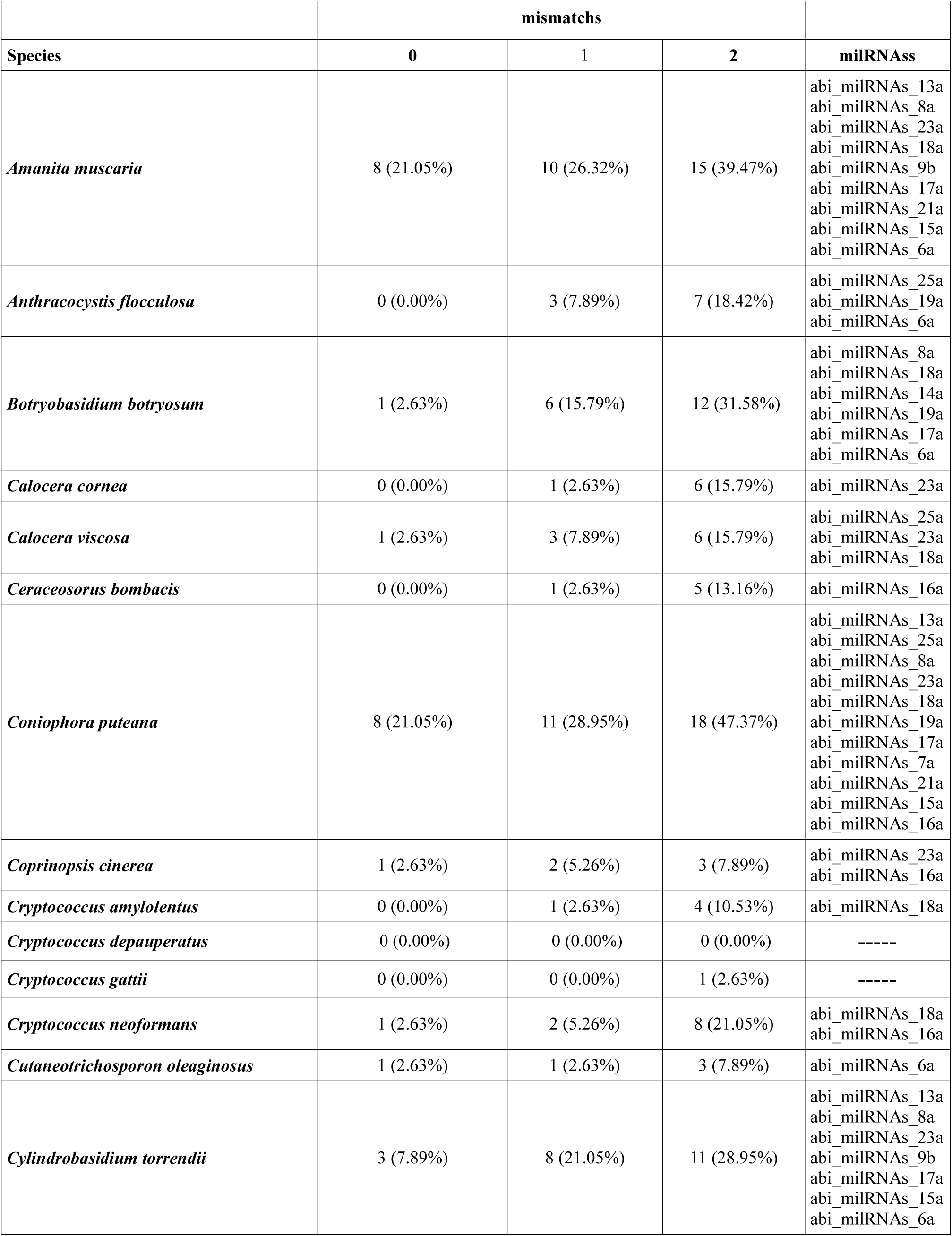
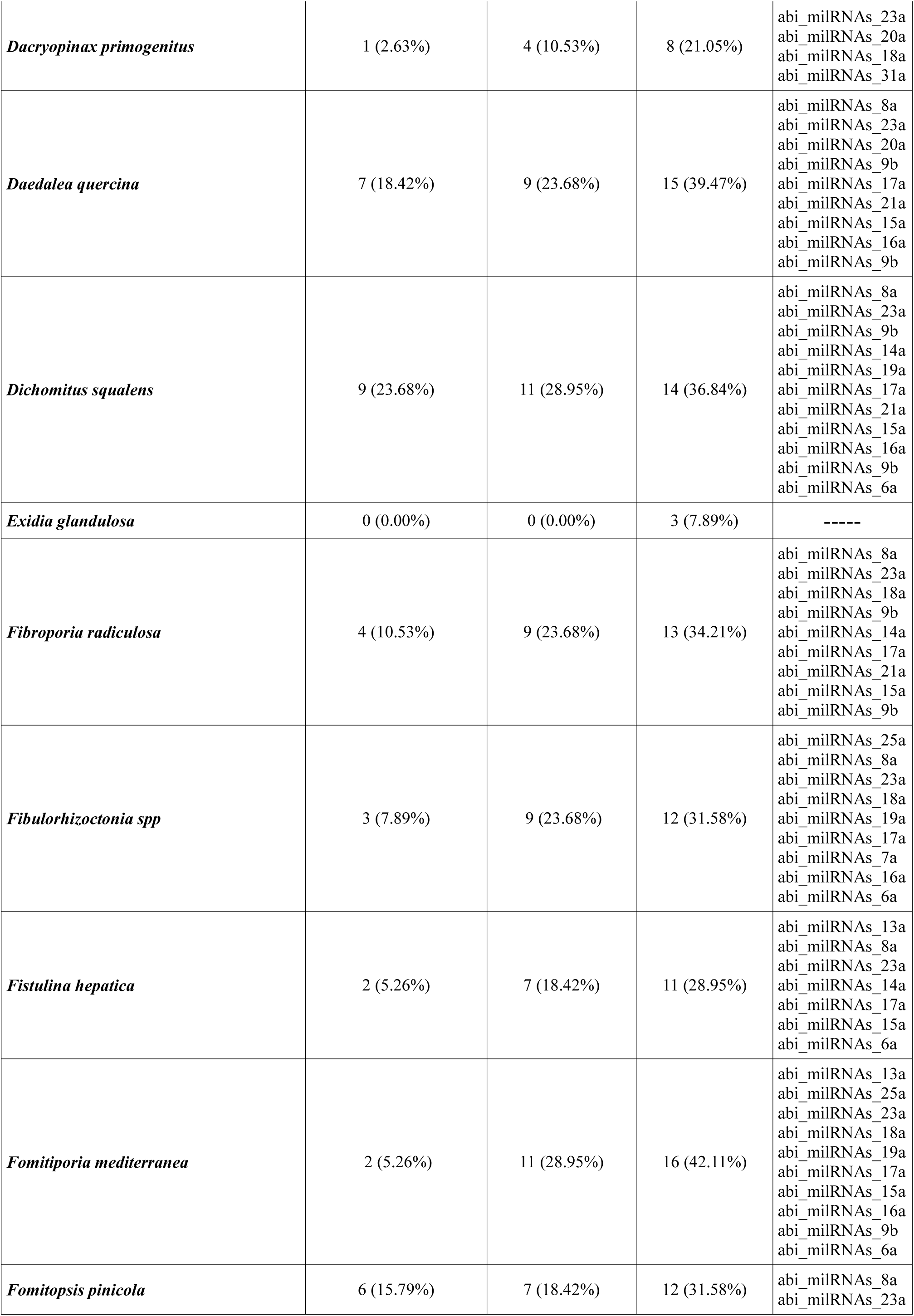
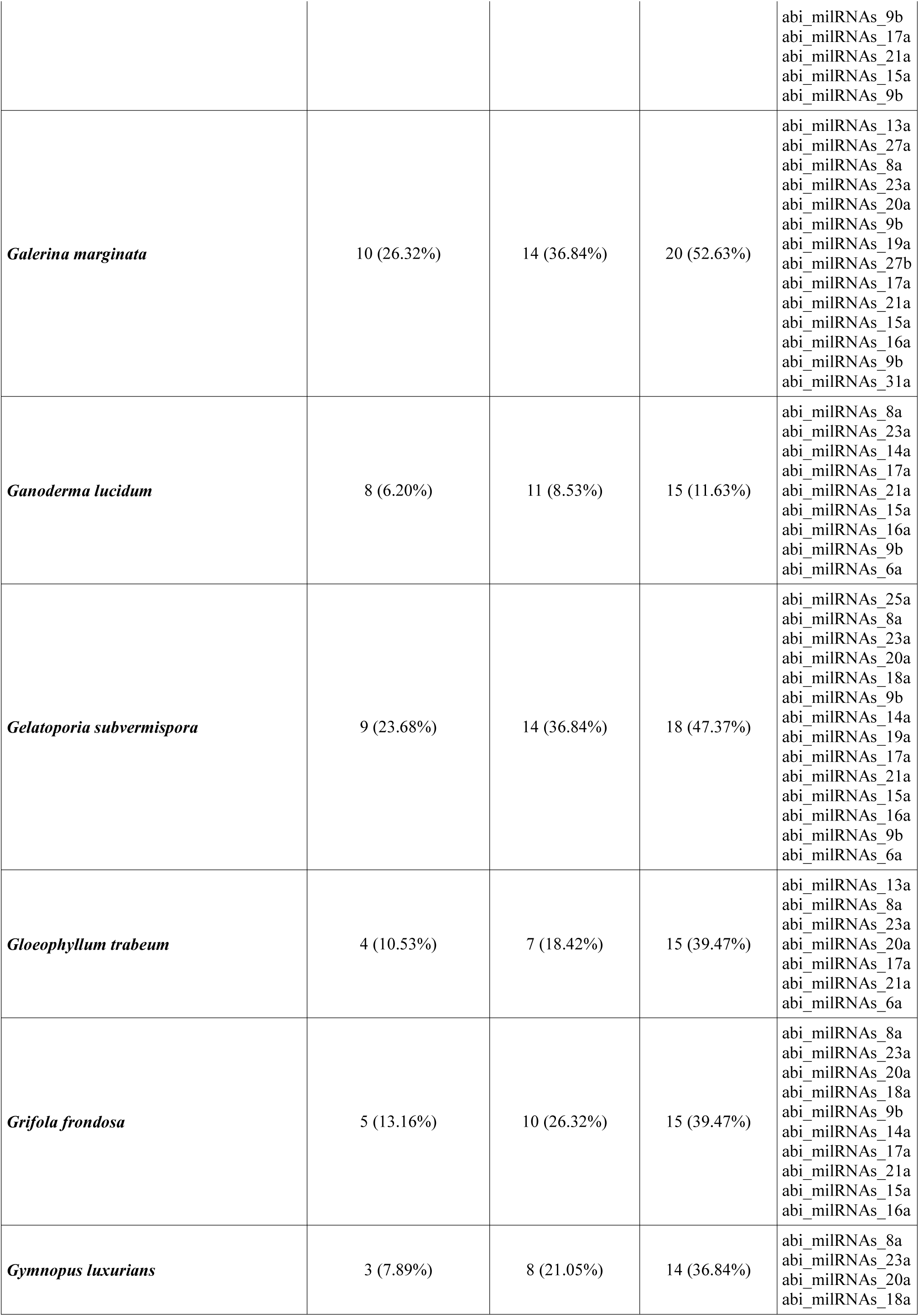
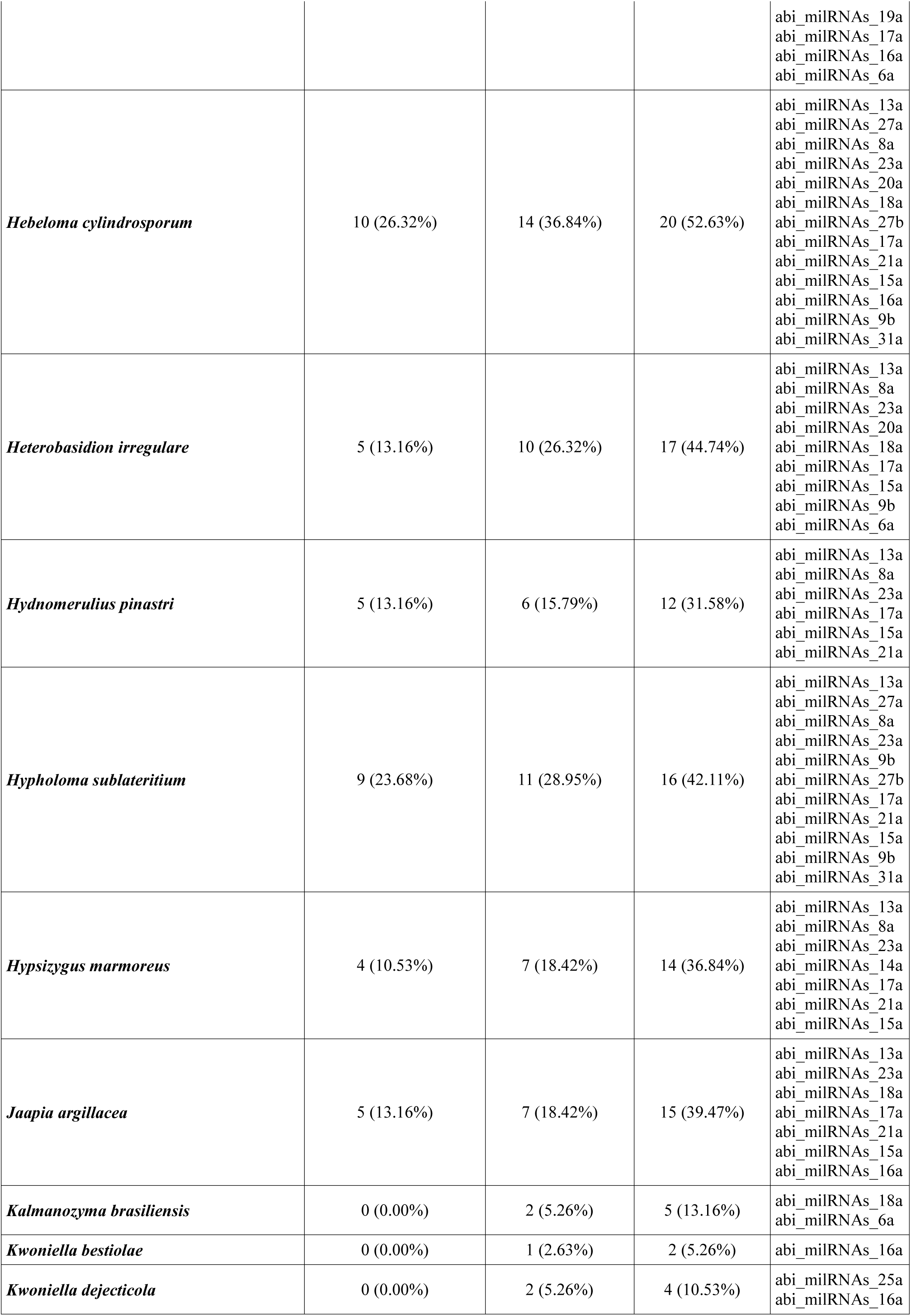
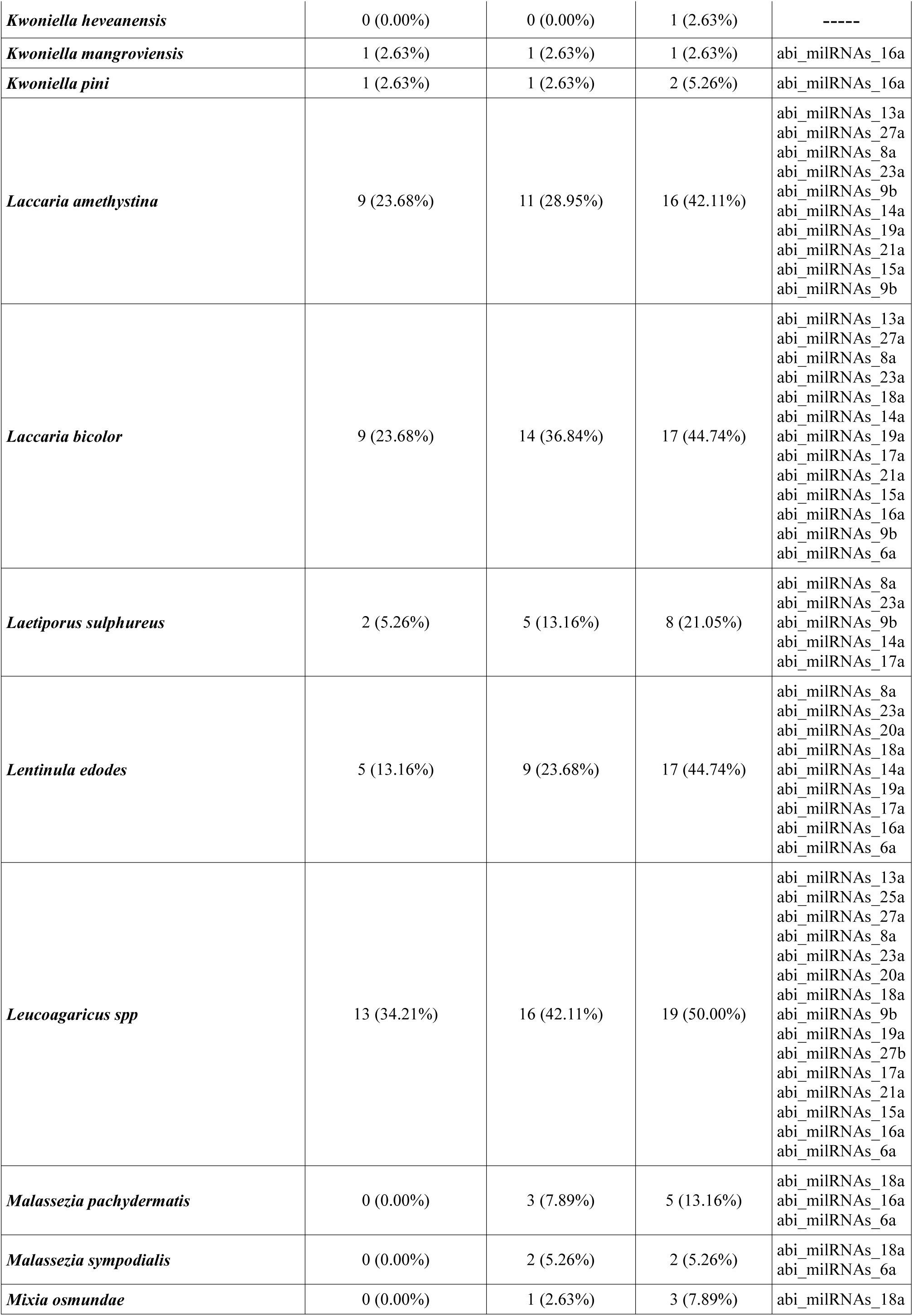
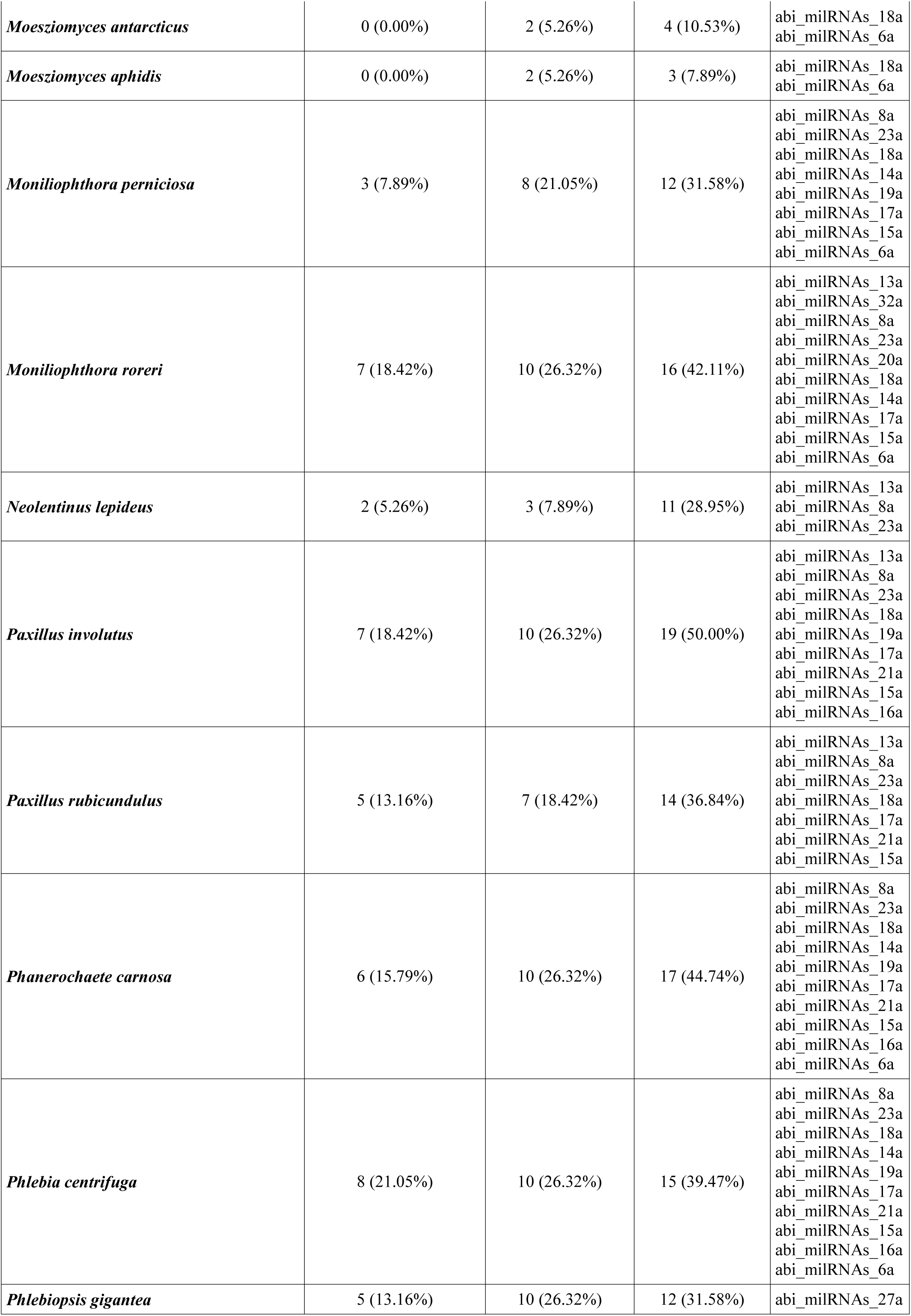
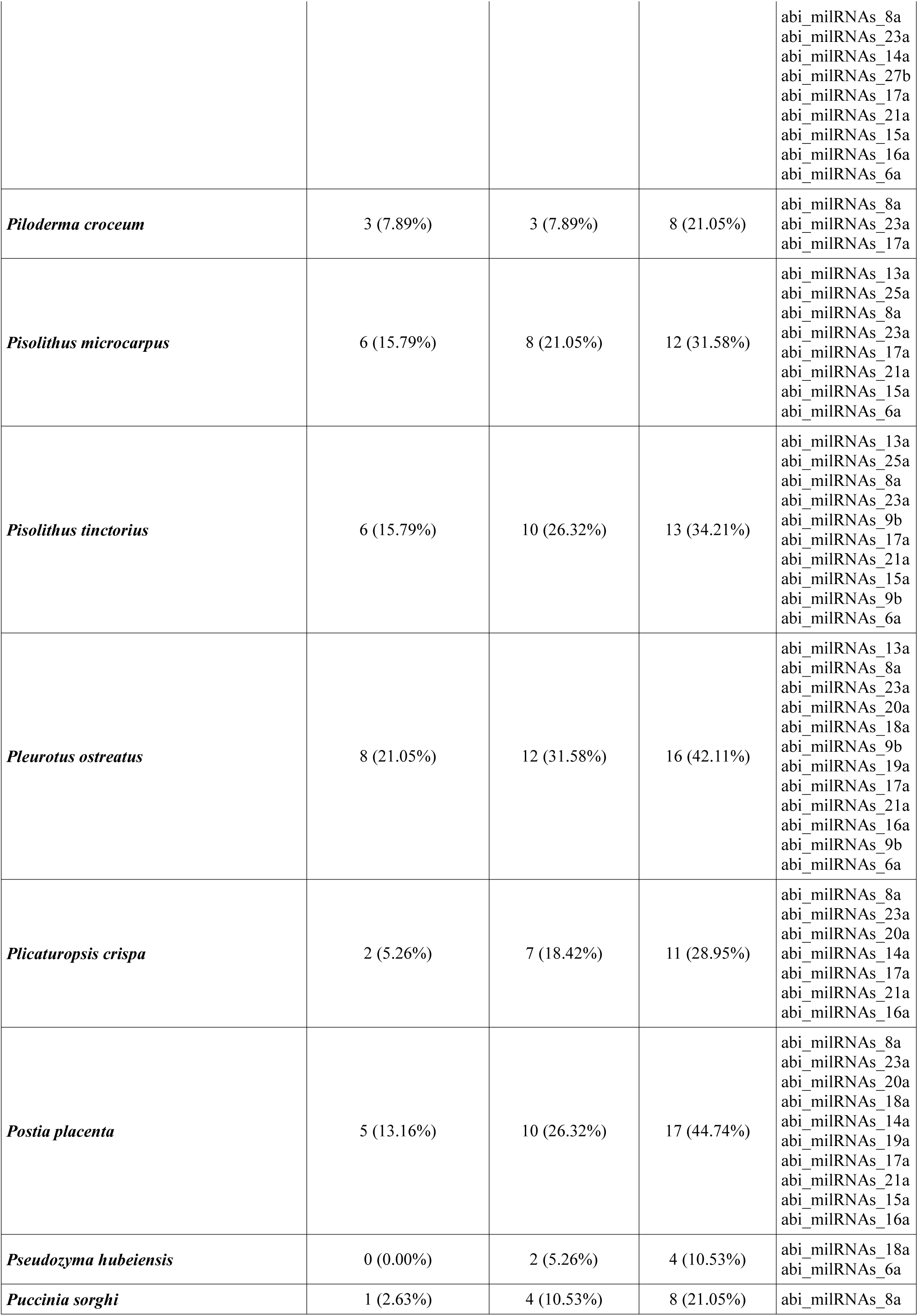
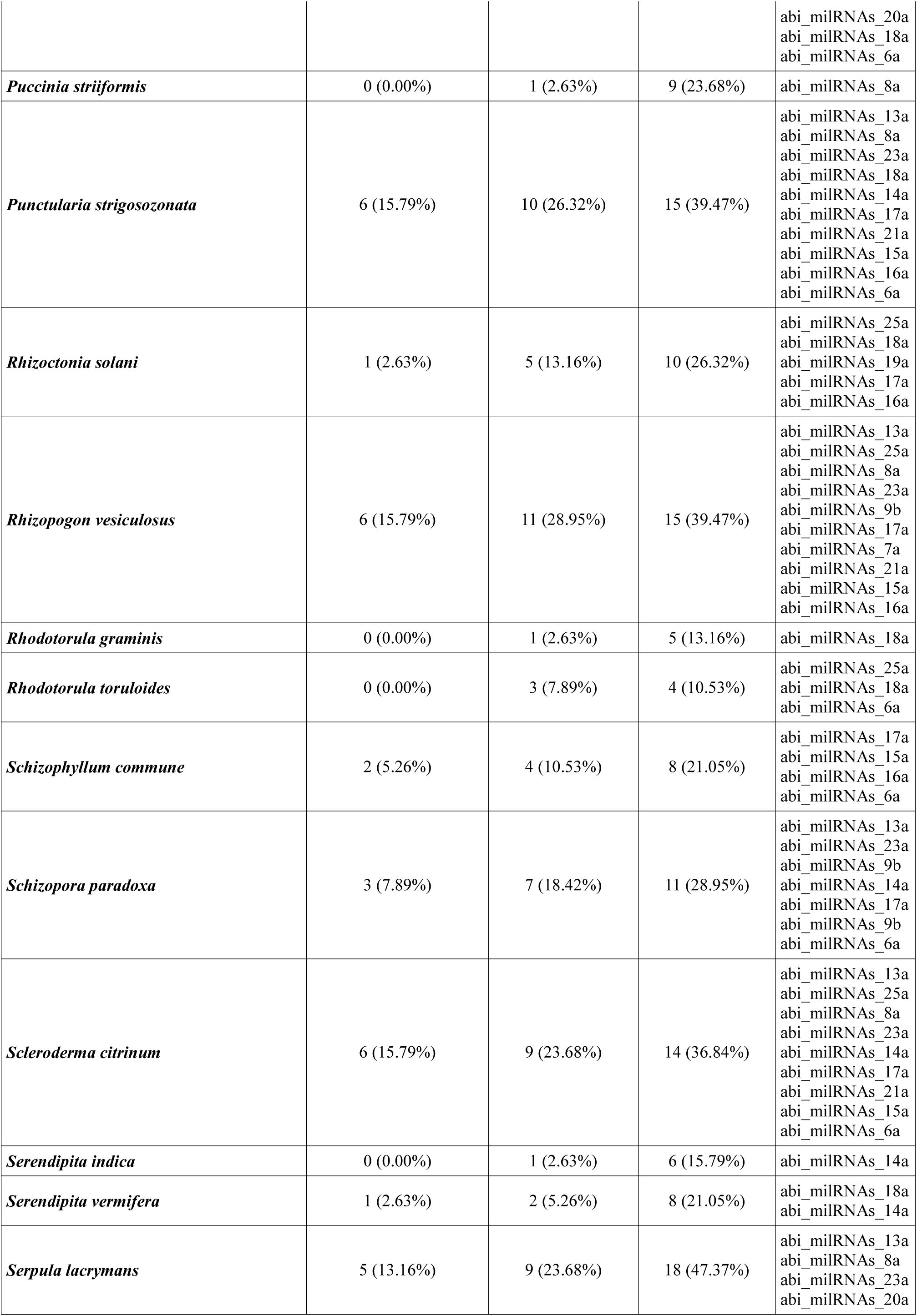
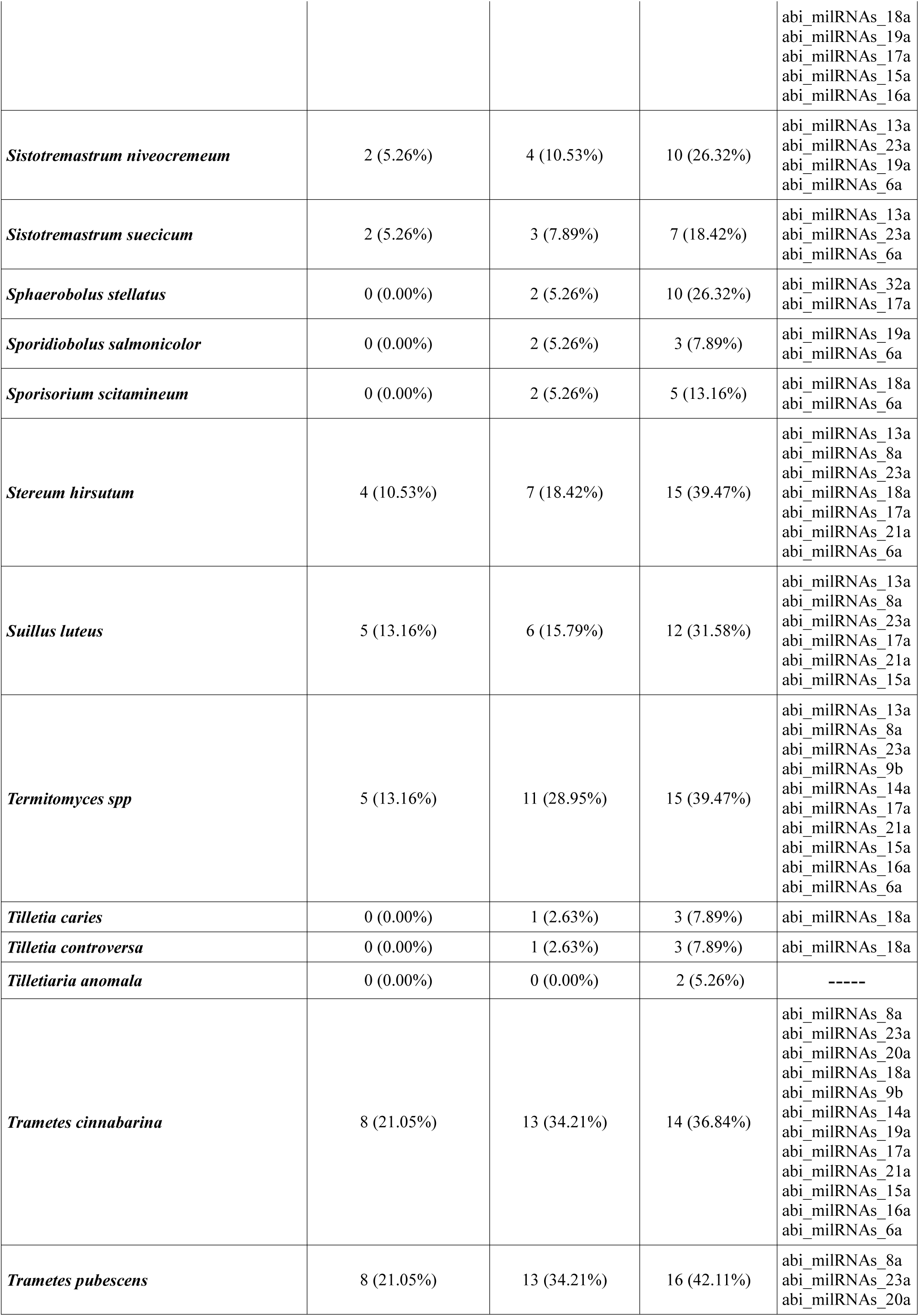
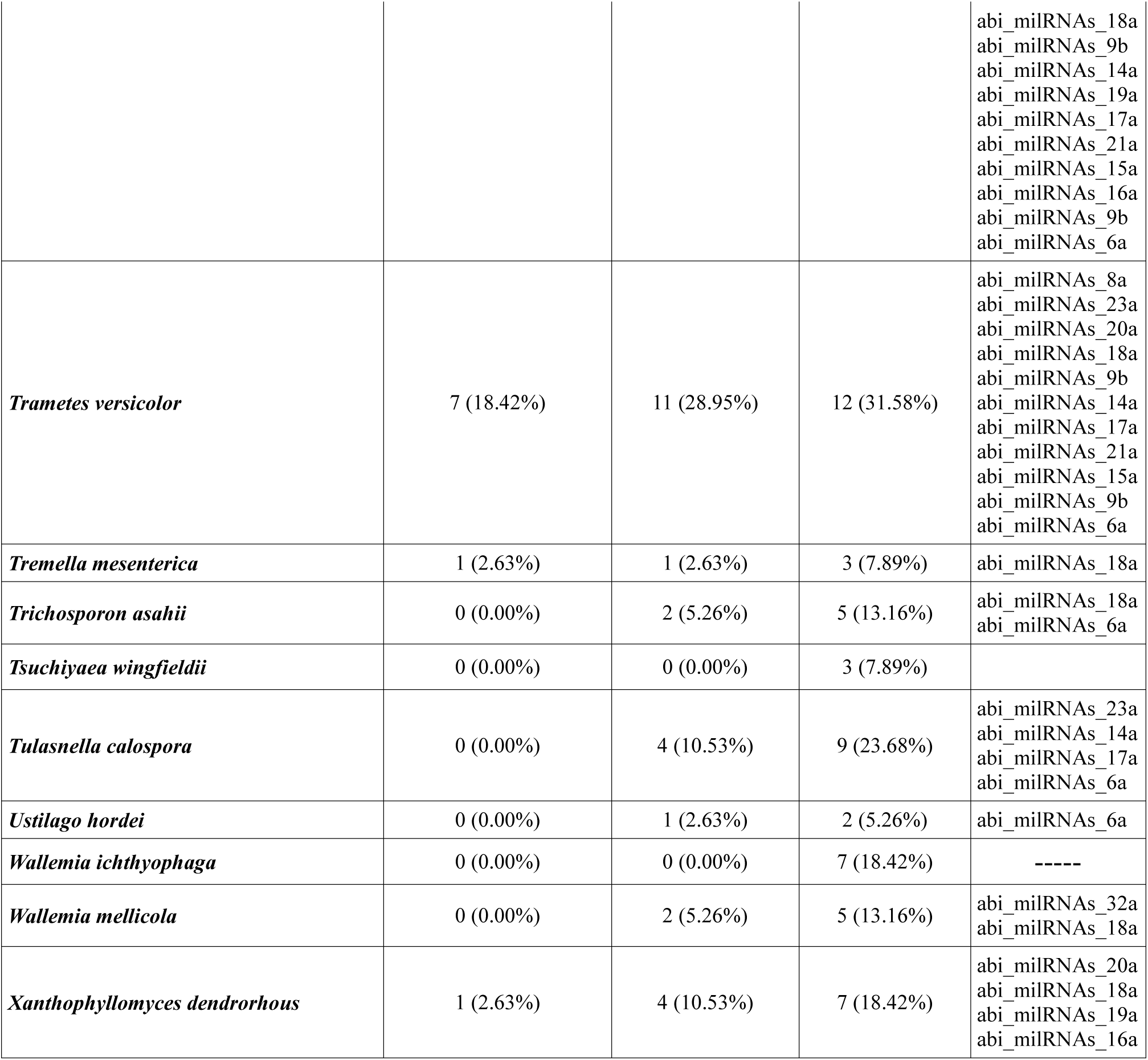
Homology between 37 *de novo* predicted *Agaricus bisporus* milRNAss and 104 *Basidiomycetes* species. Results show number and percentage (between brackets) for 0, 1 and 2 mismatches (from left to right, 2^nd^, 3^rd^ and 4^th^ columns). In column to the right (milRNAss) are listed those milRNAss with 0 or 1 mismatch. Species are listed in alphabetical order.

